# Repeated Viewing of a Film Clip Changes Event Timescales in The Brain

**DOI:** 10.1101/2025.08.27.672403

**Authors:** Narjes Al-Zahli, Mariam Aly, Christopher Baldassano

## Abstract

Many everyday experiences share a recurring structure: routines, familiar routes, rewatched films, and replayed songs. How do repeated encounters with such structure alter the brain’s representations of events? We hypothesized that, with repeated viewing of a film clip, event representations in the brain may adapt by becoming either finer (more detailed) or coarser (more generalized). To test this hypothesis, we analyzed data from 30 human participants (12 males, 18 females) who underwent functional magnetic resonance imaging (fMRI) while watching three 90-second clips from “The Grand Budapest Hotel” six times each. We used hidden Markov models and pattern similarity analysis applied to searchlights across the brain to quantify the strength of event structure at different timescales for each clip presentation. We then tested how event structure strength changed at both slow and fast timescales with repeated viewings. Most brain regions exhibited stability in the strength of event structure at both slow and fast timescales. Other regions, however, showed flexible event representations that became more or less granular across repeated clip presentations. Notably, several brain regions exhibited consistent changes in the strength of event structure at a slow timescale across different movie clips. Furthermore, in lateral occipital cortex and middle temporal gyrus, slow timescale structure was correlated with subsequent memory for the clips. These results highlight that event dynamics in the brain are not fixed, but can change flexibly with experience.

**Significance Statement:** Many day-to-day experiences recur over time, as we retrace the same route to work or listen to our favorite song on repeat. We asked how increasing familiarity with an experience changes the brain’s representations of it. Individuals repeatedly watched film clips while undergoing fMRI. We examined how the brain’s temporal representations of events in the clips changed with repeated viewing. As clips became familiar, some brain regions exhibited fine-tuned event representations that divided film clips into smaller events. Other brain regions showed coarser event representations that grouped previously distinct events. The strength of event structure at a coarse timescale was correlated with memory. These results show that the brain flexibly changes how it represents events as they become more familiar.

## Introduction

Experience unfolds over multiple timescales—from the millisecond flux of sensations to minutes-long narratives. Neuroimaging, electrophysiological, and computational work shows that the brain mirrors this temporal structure by processing information hierarchically across multiple timescales. Lower-level sensory cortex operates on short timescales, rapidly tracking moment-to-moment changes in sensory input, whereas higher-order association areas accumulate information over progressively longer periods. This gradient of “temporal receptive windows” (<u>Hasson et al., 2008</u>; <u>Honey et al., 2012</u>; <u>Murray et al., 2014</u>; <u>Geerligs et al., 2022</u>) supports segmentation of continuous experience into discrete events at multiple timescales, which in turn scaffolds situation models and episodic memories that weave discrete moments into narratives (<u>Zacks et al., 2001</u>; <u>Baldassano et al., 2017</u>).

While research has established different timescales of processing across the brain, it remains an open question how flexible these timescales are. A given region’s temporal integration window may be determined by local circuit properties and its position in the cortical hierarchy (<u>Chaudhuri et al., 2015</u>; <u>Hasson et al., 2008</u>), indicating that a brain region’s processing timescale may be relatively fixed. However, emerging evidence points to malleability of timescales: the brain can adapt its temporal processing to stimulus properties, context, goals, and expectations (<u>Lerner et al., 2014</u>; <u>Baumgarten et al., 2021</u>; <u>Shin & DuBrow, 2021</u>; <u>De Soares et al.,</u> <u>2024</u>; <u>Çatal et al., 2024</u>; <u>Lee et al., 2021</u>). Further, as content becomes familiar, the brain uses prior knowledge to anticipate upcoming information (<u>Favila & Aly, 2025</u>; <u>Günseli & Aly, 2020</u>; <u>Tarder Stoll et al., 2024</u>), potentially shifting the timing of event encoding (<u>Shin & DuBrow, 2021</u>; <u>Lee et al., 2021</u>; <u>De Soares et al., 2024</u>). We therefore hypothesized that repeated exposure to an event may change neural representations of event structure at multiple timescales. Repeated experience may allow the brain to group sub-events that were previously perceived to be separate, or fine-tune larger event structures into subcomponents. These changes may be detectable as speeding or slowing of the brain’s event representations with repeated experience, even if the external stimulus remains unchanged.

Here, we examined such experience-driven shifts in event timescales in the brain. Characterizing event timescales can be complex: even within a single brain region, multiple timescales of activity can be hierarchically nested (shorter events embedded within longer events) (<u>Stephens et al., 2013</u>; <u>Kanter et al., 2025</u>). To detect event structure at multiple concurrent timescales within individual brain regions, we developed a method that first segments continuous neural activity into discrete events at multiple timescales, and then quantifies the strength of event structure at each timescale. We applied this method to fMRI data from <u>Aly et al.</u> <u>(2018)</u>, in which participants repeatedly viewed the same movie clips (**Fig. 1**). The clips varied in narrative coherence (continuous storyline vs. scrambled scene order), and temporal stability (fixed vs. variable scene order across repetitions), allowing us to examine how neural event structure changes across different story structures with different opportunities for learning shorter-timescale events, longer timescale regularities, and the overall narrative. Narrative coherence determines whether meaningful long-timescale structure is available to the viewer on the first exposure: a coherent storyline (Intact clip) enables the immediate extraction of overarching narrative gist, whereas scrambled narratives (Scrambled-Fixed and Scrambled-Random clips) disrupt causal and thematic continuity, limiting the ability to understand slowly unfolding narratives on initial viewing. Temporal stability, on the other hand, determines how much event structure is available to be learned across repetitions. When the order of scenes is fixed (Intact; Scrambled-Fixed), viewers can gradually learn transitional probabilities between scenes and develop expectations about upcoming scenes even in the absence of a coherent narrative; when the scene order changes each viewing, such learning is impossible (Scrambled-Random). We therefore studied conditions that differ in the degree to which slow-timescale (narrative-level) and fast-timescale (scene transitions) event structure is immediately available or can be learned with experience. This allowed us to test whether familiarity-driven changes in neural event timescales depend on the presence of meaningful narrative structure, the possibility of learning temporal regularities, or both.

**Figure 1.**
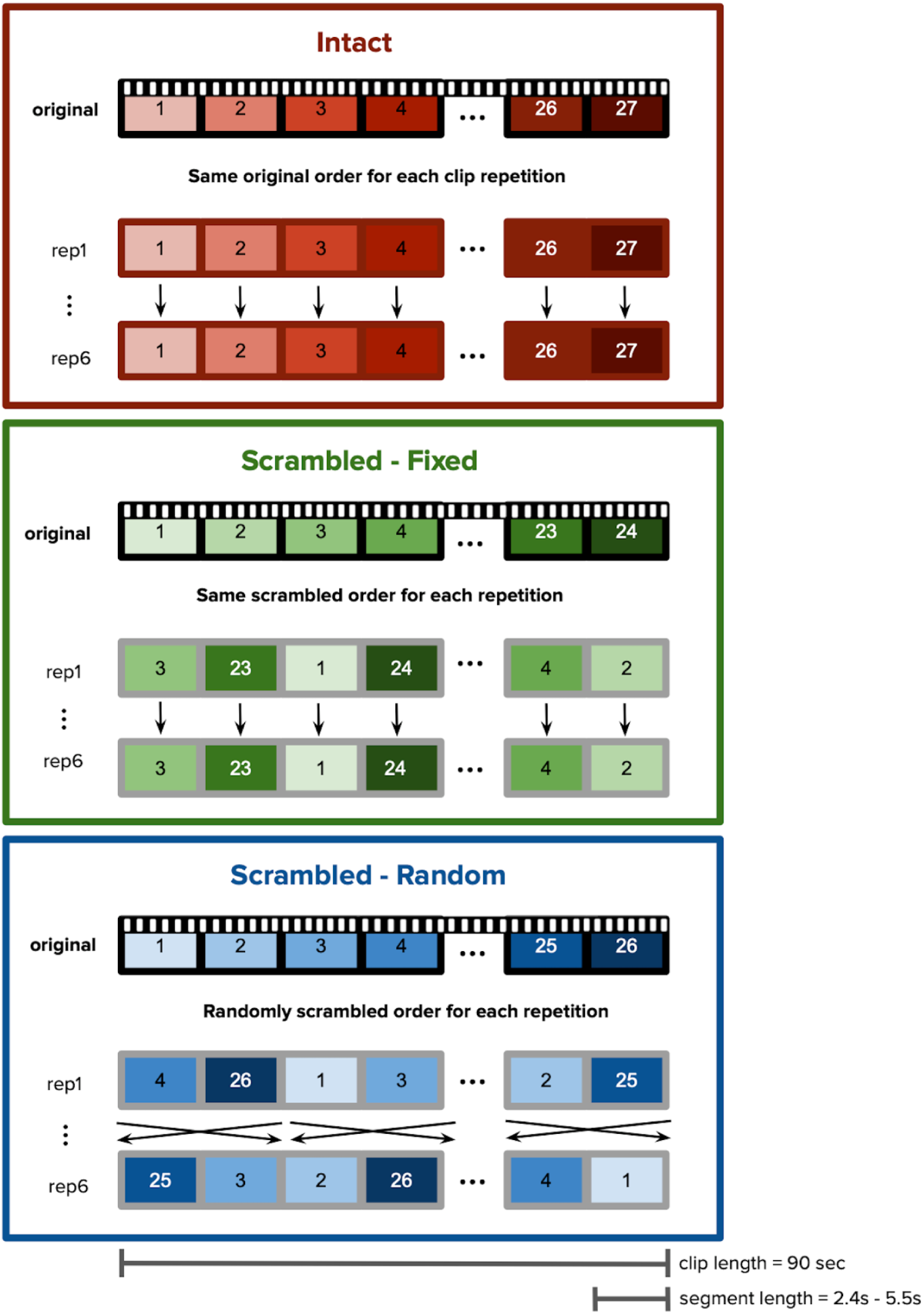
Stimuli and Experimental Design. Participants watched three 90-second clips from *The Grand Budapest Hotel*. Each clip was watched six times. Each clip was segmented into smaller continuous chunks, 2.4-5.5s long. The numbers in each clip refer to a segment within each clip. Each clip was assigned to one of three conditions. In the *Intact* condition, the clip segments were shown in their original order. In the *Scrambled-Fixed* condition, the clip segments were randomly shuffled, but shown in the same scrambled sequence for all clip presentations. In the *Scrambled-Random* condition, the clip segments were randomly shuffled and shown in a different order for each clip presentation.

To address these questions, we used a Hidden Markov Model-based event segmentation approach (<u>Baldassano et al., 2017</u>) to identify event boundaries in each brain region across multiple timescales, and used pattern similarity to examine the strength of event representations within and across event boundaries. This approach, applied to each movie viewing separately, allowed us to: (i) determine how event structure at slow and fast timescales changed with experience; (ii) assess which changes in event structure were specific to a particular movie clip vs. generalized across clips with different content and temporal structure; and (iii) relate these changes to memory. Together, this approach tested flexibility vs. stability in the timescales of event processing in the brain, and the relevance of these changes for memory.

## Materials and Methods

### Experimental Design

#### Grand Budapest Hotel dataset

We used data collected by <u>Aly et al. (2018)</u>. The dataset included thirty right-handed participants (12 men, average age = 23.0 years, SD = 4.2; average education = 15.3 years, SD = 3.2). During functional magnetic resonance imaging (fMRI), these participants watched three 90-second clips from *The Grand Budapest Hotel* (**Fig. 1**). Each clip was shown six times. The clips were pseudo-randomly interleaved, such that all three were shown one time each before the next set of presentations began. None of the participants had seen the movie before. After viewing the three clips six times each, participants were asked to recall the clips. They were given a blank text document and instructed to type their memories for the three clips in any order they chose.

The study included three different types of clips. Each 90-second clip was divided into smaller segments, 2.4-5.5 seconds in length. The first clip, *Intact*, was shown in its original narratively coherent order, i.e., the segments were viewed in the same order as the original movie. The second clip, *Scrambled-Fixed*, featured randomly ordered segments, but the order was consistent across viewings. This resulted in a stable but incoherent structure. The last clip, *Scrambled-Random*, featured segments that were reordered differently each time, creating an incoherent and unstable structure.

Three 90-second movie scenes were selected: an interview scene, a painting theft scene, and a chase scene. There were two counterbalancing groups, in which the assignment of movie scene to condition (*Intact, Scrambled-Fixed, Scrambled-Random*) was partially counterbalanced.

For both groups, the *Intact* clip was the interview scene. The *Scrambled-Fixed* clip was the painting theft scene for one group and the chase scene for the other group, and vice versa for the *Scrambled-Random* clip.

#### fMRI Methods

##### fMRI Acquisition

Full details of the MRI acquisition and preprocessing procedures are reported in <u>Aly et al.</u> <u>(2018)</u>. Functional MRI data were acquired using a 3-Tesla Siemens Prisma scanner equipped with a 64-channel head/neck coil. Functional images were collected using a multiband echo-planar imaging (EPI) sequence with the following parameters: repetition time (TR) = 1.5 s, echo time (TE) = 39 ms, flip angle = 50°, multiband acceleration factor = 4, and slice shift = 3. Each volume consisted of 60 oblique axial slices acquired in an interleaved order, with an isotropic voxel size of 2.0 mm. High-resolution T1-weighted structural images were obtained with an MPRAGE sequence (1.0 mm isotropic resolution) for anatomical reference and registration. Additionally, field maps were collected for registration (40 oblique axial slices, 3.0 mm isotropic resolution). The fMRI scanning session consisted of three experimental runs, during which participants viewed clips from the Intact, SFix, and SRnd conditions.

##### fMRI Preprocessing

Data preprocessing and registration were performed using tools from FEAT, FLIRT, and command-line functions in FSL (http://fsl.fmrib.ox.ac.uk/fsl/). The first three EPI volumes of each run were discarded to allow for T1 signal equilibration. Preprocessing steps included brain extraction, motion correction, high-pass temporal filtering (cutoff period = 140 s), and spatial smoothing (3-mm FWHM Gaussian kernel). Field maps were processed according to the FSL FUGUE user guide; magnitude images were averaged and skull-stripped, while phase images were converted to rad/s and smoothed (with a 2-mm Gaussian kernel). The resulting phase and magnitude images were used to unwrap the functional images in the preprocessing step of FEAT analysis to reduce distortions and aid registration to anatomical space. Functional data were then registered to each participant’s anatomical scan, followed by normalization to standard Montreal Neurological Institute (MNI) space. After preprocessing, functional time series for each run were divided into volumes according to individual movie clip viewings, with the initial four volumes of each clip viewing removed to account for hemodynamic lag. Finally, the timeseries for each clip was z-scored within each participant to ensure comparability across runs and participants.

### fMRI Visualization

Functional data were projected from volume space to cortical surface space using Nilearn for visualization purposes only; all analyses were conducted in volume space using custom Python scripts and libraries.

### Whole-Brain Searchlight Analysis

We conducted a whole-brain analysis using spherical searchlights generated by <u>Lee et al.</u> <u>(2021)</u>. The searchlights were spaced evenly throughout the MNI volume (radius = 5 voxels; stride = 5 voxels). We only analyzed searchlights containing at least 20 voxels within a standard MNI brain mask and with valid data from at least 15 participants for all viewings. For each voxel, we computed a Within- vs. Between-Event Similarity value (defined in the Statistical Analysis section below) by averaging the Within- vs. Between-Event Similarity values of all searchlights containing that voxel, separately for each of the 6 viewings for each of the 3 clips.

### Statistical Analysis

#### Scoring of event memory

Free recall data were scored by counting the number of details reported by each participant for each movie clip. Details included characters, dialogue, actions, and perceptual details such as the appearance of characters and the spaces through which they moved. Reported details were checked against the movie clips for veracity. Points were only given for accurate details.

#### Filtering for Regions with Meaningful Timescales Across Clip Viewings

##### Event Segmentation Model

We used the event segmentation Hidden Markov Model (HMM) described by <u>Baldassano</u> <u>et al., 2017</u> to identify event boundaries for each searchlight in the brain (**Fig. 2**). The HMM operates under the following assumptions: (1) a brain region’s response to a dynamic stimulus consists of a sequence of discrete event states, and (2) each event is represented in a brain region by a unique spatial activity pattern. We fit the HMM to each searchlight in the brain, independently for each presentation of each clip, varying the number of events from 2 to 10 to capture event structure at slower timescales (few events) and faster timescales (more events). This resulted in nine models for each of the six viewings for each of the three clips. These event numbers covered the range of plausible event timescales for meaningful events in the video (from ∼8 seconds to 45 seconds). Each model provided the location of event boundaries for the specified number of events (2-10 across models) for each searchlight and each viewing of each clip. These event boundaries were used in subsequent analyses, described below. This approach deviates from prior work in that we do not summarize each brain region’s activity with a single timescale, nor do we assume that the voxel pattern representing each event remains stable across movie repetitions.

**Figure 2.**
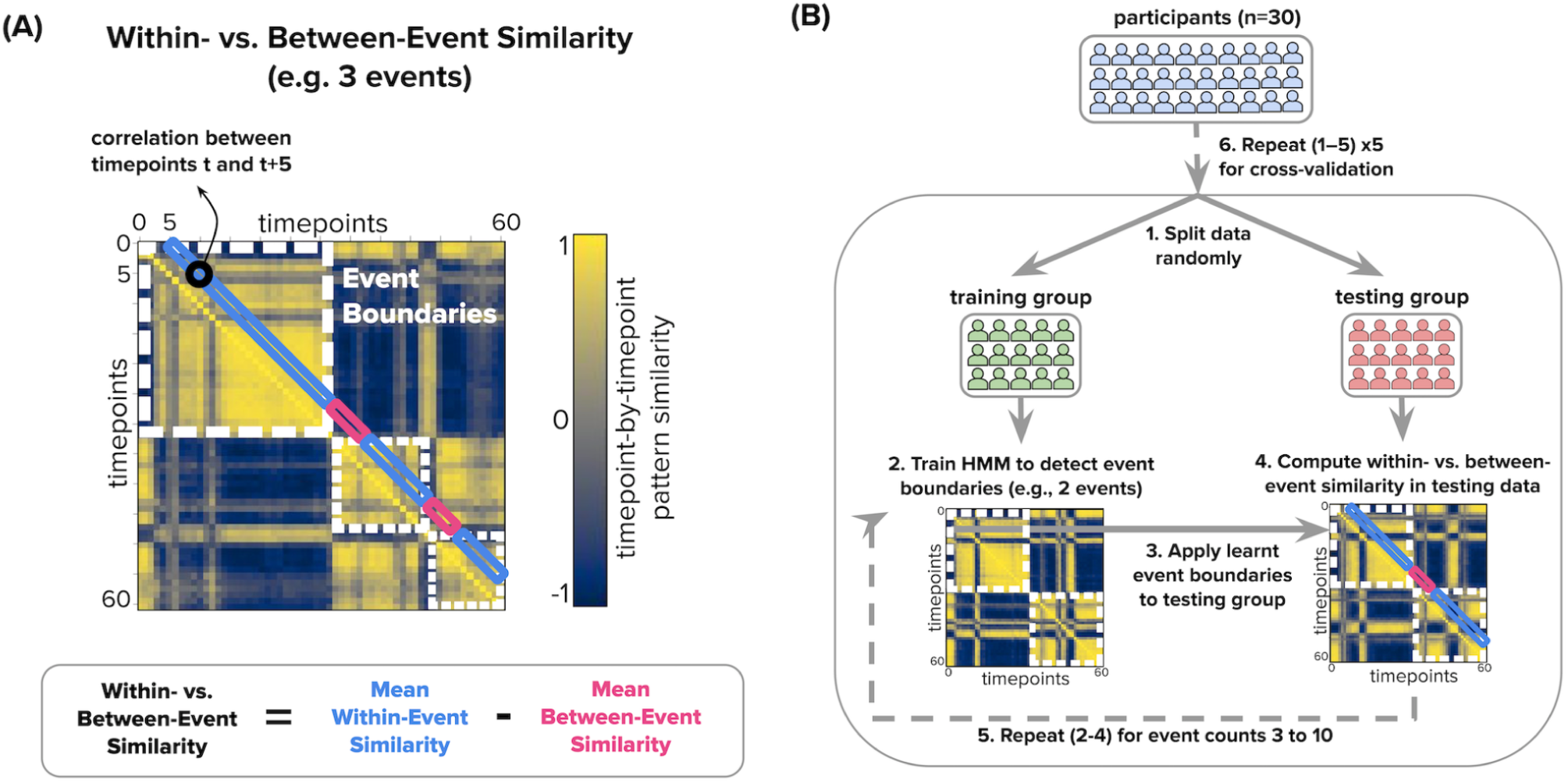
Computing Within- vs. Between-Event Similarity for Each Clip Presentation. We used a Hidden Markov Model (HMM) to identify event boundaries in each clip presentation and then assessed Within-Event vs. Between-Event pattern similarity in an independent group of participants. The analysis pipeline is visualized here with 3 events, but the model was fit for 2-10 events to assess event structure at multiple timescales. **(A)** The analysis is based on identifying stable patterns of activity within each searchlight in the brain, which correspond to a neural “event”. For each clip presentation, a Timepoint-by-Timepoint Similarity Matrix visualizes the correlation between the spatial patterns of activity for two timepoints in the clip. Dashed white outlines indicate the event boundaries identified by the HMM when the model segments the neural data into 3 events. The strength of these model-identified event boundaries was measured by computing the difference in pattern similarity for pairs of timepoints within the same HMM-defined event (blue rectangles) versus between events (red rectangles). To minimize the effects of BOLD autocorrelation, we examine Within-Event and Between-Event correlations at a delay of 5 TRs. **(B)** To produce an unbiased estimate of Within- vs. Between-Event Similarity, data from participants were randomly split into training and testing groups. Averaged time series data from the training group were used to identify event boundaries with the HMM (here, 3 events; the process is conducted for event counts 2-10). Within- vs. Between-Event Similarity around these training-group HMM boundaries was calculated for neural patterns in the testing group and used for subsequent analyses. This approach is repeated 5 times, with different participant splits.

### Within- vs. Between-Event Similarity

To evaluate the quality of event segmentation, we computed the similarity of spatial activity patterns within versus across HMM-defined event boundaries, using only held-out (testing) data. Event boundaries were defined by fitting the HMM (described above) to the group-averaged BOLD activity timecourse from a randomly selected half of the participants and identifying the timepoints at which there was a change in the most-probable event label. These boundaries were then applied to the group-averaged BOLD activity timecourse from the held-out participants. For each clip, we generated a timepoint-by-timepoint similarity matrix showing the correlation between brain activity patterns across time. To minimize the influence of BOLD autocorrelation, we calculated correlations at a fixed lag of 5 TRs. Within-event correlations were defined as pairs of timepoints separated by this lag that fell within the same event, while between-event correlations spanned an event boundary. The Within- vs. Between-Event Similarity value was calculated as the difference between the average within-event and between-event correlations, providing a measure of how well the event structure learned from training data generalized to stable neural activity patterns in unseen data. Values above zero indicate that patterns are more similar within events than between events (i.e., indicate detectable event structure at that event count).

The training and testing sets were then swapped, and the Within- vs. Between-Event Similarity values from both directions were averaged. To avoid overfitting and ensure robustness, we averaged Within- vs. Between-Event Similarity values from five different participant splits for each model. Because the movie clips used in the Scrambled-Fixed and Scrambled-Random conditions were counterbalanced across participants, the training/testing split for these clips was done within each counterbalancing group separately, and the Within- vs. Between-Event Similarity measures were averaged across the counterbalancing groups. For each clip, this procedure produced 54 Within- vs. Between-Event Similarity values (nine event counts for each of six clip presentations).

To validate the reliability of the Within- vs. Between-Event Similarity metric across participants, we measured the stability of this metric across subsets of the Intact dataset. Specifically, we randomly divided participants into two independent datasets. Within each of these datasets, we carried out the procedure described above: We identified HMM event boundaries in half of the participants (training group) and then used those boundaries to compute Within- vs. Between-Event Similarity in the other half of participants (testing group). As before, training and testing sets were then swapped, and results were averaged across five different participant splits. We repeated this procedure across event counts, viewings, and searchlights (limited to those showing meaningful event structure across all viewings; see *Statistical Significance Testing*, below). Finally, we examined between-group reliability in the Within- vs. Between-Event Similarity values. High agreement (mean Pearson r = 0.48, range = 0.36–0.59 for initial viewing; mean Pearson r = 0.59, range = 0.42–0.69 for subsequent viewings) between the two independently derived Within- vs. Between-Event Similarity values demonstrated that the measure is a stable measure across subsets of participants (**Figs. S1 and S2**).

### Statistical Significance Testing

We tested whether Within- vs. Between-Event Similarity values were statistically greater than zero using a permutation-based null hypothesis approach. Null event boundaries were generated by randomly shuffling the order of the event segment lengths from the trained HMM model, producing boundaries that had the same spacings as in the real analysis but were placed on arbitrary timepoints. Within- vs. Between-Event Similarity values for these null boundaries were then computed on the testing data. Importantly, permutations were constrained such that the shuffled sequence never matched the real sequence of event neural patterns to ensure a valid null comparison. For each event count, we conducted up to 50 permutations, calculating the Within- vs. Between Event Similarity value for each permutation. Note that for event counts below 5, there were fewer than 50 possible permutations, and so we instead used all possible shuffled permutations. We then fit a normal distribution to all of these values (across event counts) to create a single null distribution for each searchlight, clip, and viewing. The p-value was then calculated by computing the area under the normal distribution that surpassed the real Within- vs. Between-Event Similarity for each event count. This pooled approach is appropriate because the per-event-count null distributions of the Within- vs. Between-Event Similarity values are centered at the same value (≈0) and show comparable dispersion, making samples effectively exchangeable (**Fig. S3**). Pooling therefore allows us to compute p-values for all event counts in a consistent way.

For subsequent analyses, we only kept searchlights for which, for each clip, every viewing had at least one event count that produced a Within- vs. Between-Event Similarity value above zero and significantly greater than the null distribution (p > 0.05, one-tailed). This ensured that only searchlights with reliable event structures for all viewings proceeded, enabling subsequent identification of changes in event structure.

#### Identifying Changes in Event Structure

We quantified changes in event structure across repetitions of each clip at both slow (2-event) and fast (10-event) timescales. Event structure strength is defined as the Within- vs. Between-Event Similarity for a given event count; higher values indicate more distinct event representations. Change was defined as the difference between the average Within- vs. Between-Event Similarity across repeated viewings (viewings 2–6) vs. the first viewing, separately for 2 events and 10 events (**Fig. 3**). Examples of these changes are illustrated in timepoint-by-timepoint similarity matrices (Figs. S4–S5). We focused our statistical comparisons on the first viewing (no prior familiarity with the clips) vs. subsequent viewings (varying levels of familiarity with the clips) because our primary hypotheses were about the effects of familiarity generally rather than a specific rate of change as a function of the amount of familiarity. We nevertheless visualize results for each clip viewing separately for interested readers. Likewise, we limited our statistical analyses to the slowest (2-event) and fastest (10-event) timescales that we examined, to reduce the number of statistical comparisons and convey results in a concise manner. Results are nevertheless visualized across all event counts for transparency.

**Figure 3.**
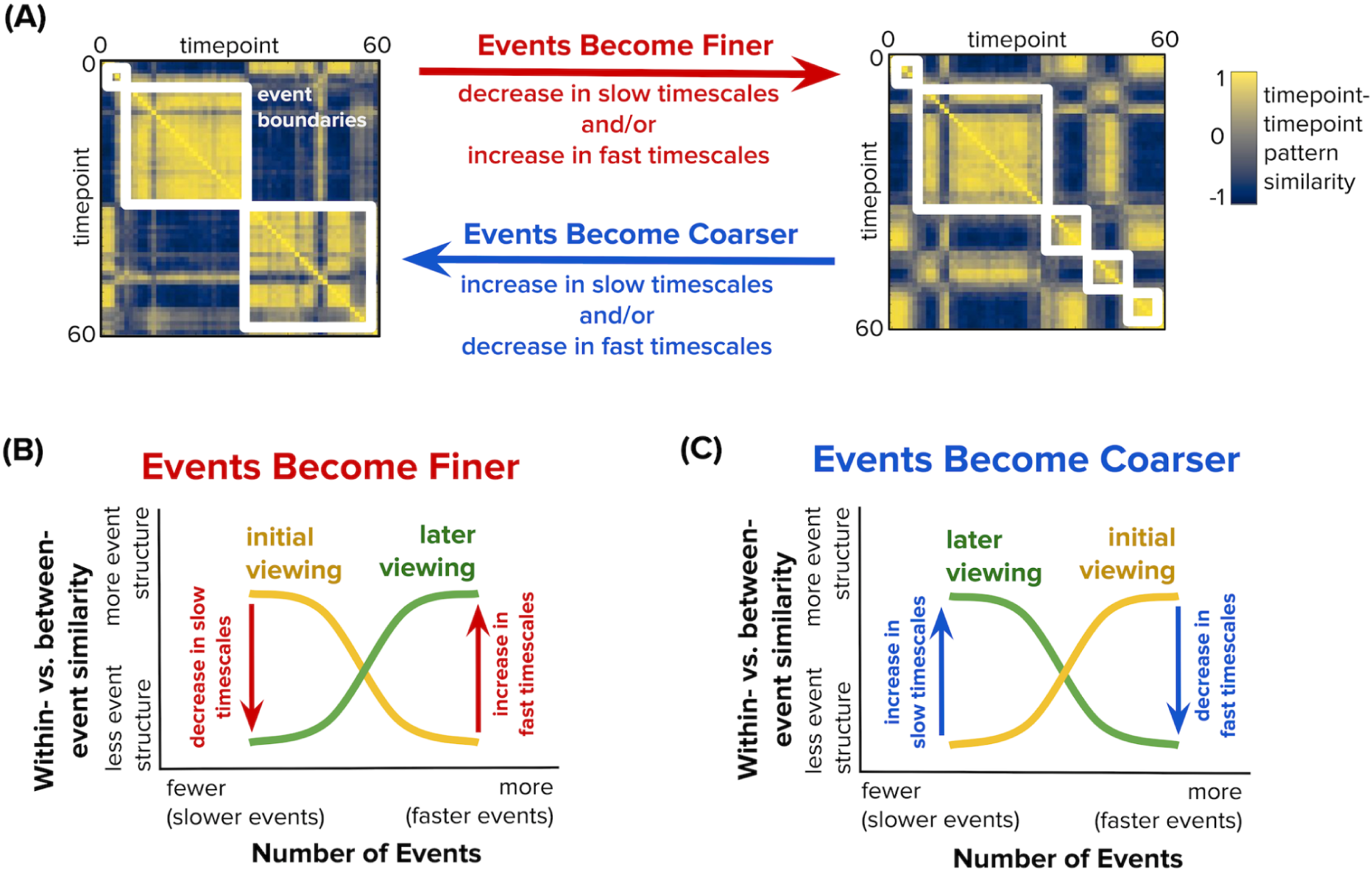
Computing Changes in Event Structure at Slow and Fast Timescales Across Clip Presentations. **(A)** Example timepoint-by-timepoint pattern similarity matrices with event structure at different timescales. The white boxes overlaid on each matrix denote event boundaries identified by the HMM analysis, capturing clusters of timepoints that are highly similar to each other. The left matrix shows a relatively coarse event segmentation, as indicated by a small number of large white boxes (events). The right matrix shows a relatively fine event segmentation, as indicated by a larger number of small white boxes (events). A brain region’s event segmentation may change with clip repetition, becoming either finer or coarser. Event representations become finer when a region exhibits weaker event structure at slow timescales over clip repetition, stronger event structure at fast timescales over clip repetition, or both—leading to more, shorter events. Conversely, event representations become coarser when a region exhibits stronger event structure at slow timescales over clip repetition, weaker event structure at fast timescales over clip repetition, or both—resulting in fewer, longer events. **(B)** We fit HMMs to each presentation of each clip, measuring the strength of event structure for each viewing (within- minus between-event similarity) while varying the number of events in the models. In this example, the initial clip presentation (yellow) shows event structure at slow timescales (fewer events), whereas subsequent clip presentations (green) show event structure at faster timescales (more events). With repeated viewing, events therefore become fine-tuned in this example, both because the region shows weaker event structure at slow timescales and stronger event structure at fast timescales. **(C)** Following the same conventions of (B), in this example, the initial clip presentation (yellow) shows event structure at fast timescales (more events), whereas subsequent clip presentations (green) show event structure at slow timescales (fewer events). With repeated viewing, events therefore become coarser in this example, both because the region exhibits stronger event structure at slow timescales and weaker event structure at fast timescales. Subsequent analyses focus on changes in event structure (within- vs. between-event similarity) at the lowest and highest event counts (2 and 10), although data will be visualized across the full range of event lengths.

To determine statistical significance for changes in event structure across viewings, we used a permutation-based null hypothesis testing method. Null datasets were created by randomly shuffling the order of six brain activity timecourses for each participant, corresponding to the six presentations of a given clip. In each permutation, any of the six viewings could be treated as ‘first’ (with the remaining five as ‘later’). We ran this analysis pipeline 51 times: once on the actual (unpermuted) dataset and 50 times on the null (permuted) datasets. This allowed us to examine the effect of first viewing vs. repeated viewings on the shuffled order. A two-tailed p-value was obtained by fitting a normal distribution to the null *change* in Within- vs. Between-Event Similarity values and computing the area under the normal distribution that exceeded the real *change* in Within- vs. Between-Event Similarity value for each searchlight, for each clip. Significant changes in Within- vs. Between-Event Similarity (q<0.05) in searchlights for a given clip were determined after applying the Benjamini-Hochberg FDR correction, as implemented in AFNI (<u>Cox, 1996</u>).

#### Combined Analysis of Movie Clips

We tested whether a given brain regions’ direction of change in event structure across repeated viewings were consistent across different movie clips. Specifically, we sought to identify which brain regions showed a directionally consistent change in event structure at slow (2-event) and fast (10-event) timescales with repeated exposure to a movie clip, regardless of the specific content of that movie clip. Because each clip underwent an initial filtering step (described above) to include only searchlights with meaningful event structure across all repetitions, we first identified the set of overlapping searchlights that passed this filter for all three clips. This ensured that only overlapping searchlights with reliable event structure were included in the cross-clip analysis.

Next, within this overlapping set of searchlights, we calculated the change in the Within-vs. Between-Event Similarity value from the first viewing to the average of viewings 2–6, separately for slow (2-event) and fast (10-event) timescales. This procedure was first done for each clip separately. We then grouped searchlights based on whether they showed a consistent increase or decrease in the Within- vs. Between-Event Similarity value across all three clips—meaning the direction of change was the same (stronger or weaker event structure over clip repetition) in every clip. For each clip and each type of change (stronger or weaker event structure over clip repetition) at each timescale (2-event and 10-event), we performed two-tailed statistical tests and applied FDR correction at ∛0.05. such that the conjunction of effects across all 3 clips would be FDR corrected at q < 0.05. Finally, we identified searchlights that overlapped across the FDR-corrected maps for each clip, resulting in separate maps for increases vs. decreases in the Within- vs. Between-Event Similarity value with repeated viewing, at both the slow (2-event) and fast (10-event) timescales.

#### Linking Changes in Event Structure to Memory

We examined whether changes in Within- vs. Between-Event Similarity predicted memory for the three clips. To do so, we initially focused on the searchlights from the conjunction analysis (i.e., regions that showed significant and consistent changes in event structure across all three clips). For each searchlight, we obtained each individual participant’s Within vs. Between-Event Similarity value (as described below) at the event timescale that exhibited change in event structure with repeated viewing for that searchlight at the group level. We also obtained each participant’s cross-clip average recall. Finally, we obtained the across-participant correlation between these values using a robust bootstrap linear regression model (10,000 iterations). Because this analysis involved a small set of six conjunction-defined searchlights, identified independently of the memory data, we treated it as exploratory and did not apply multiple-comparison correction. Focusing on this small set of six regions helped limit the number of brain-behavior correlations tested, and thus reduce potential multiplicity concerns. Given the restricted, exploratory scope of this analysis, we report uncorrected p-values and highlight effects with p < .05 (uncorrected).

We additionally ran exploratory, clip-specific brain-behavior correlations for each timescale (slow/fast) and direction of change (stronger/weaker with repetition). These analyses were carried out using searchlights that exhibited significant changes in event structure at the respective timescale and direction in group-level analyses. These exploratory tests encompassed a much larger number of searchlights than the conjunction-based analysis (23 brain-behavior correlations for Intact, 3 for Scrambled-Fixed and 45 for Scrambled-Random vs. the original 6 we did for the conjunction analysis); we therefore applied false discovery rate (FDR) correction across the tested searchlights to control for multiple comparisons.

To obtain individual Within vs. Between-Event Similarity values, we modified the original pipeline. For each clip, we calculated per-participant Within- vs. Between-Event similarity values by randomly splitting the participants within each counterbalancing group into two halves. The HMM model was trained on one half and tested individually on the participants in the other half, providing Within- vs. Between-Event similarity values for all event counts for each presentation for each participant.

To test whether these brain–behavior correlations reflected true change across repetitions or baseline differences at first viewing, we performed a follow-up ANCOVA. This model predicted recall from the change in Within- vs. Between-Event Similarity with repeated viewing while controlling for baseline Within- vs. Between-Event Similarity from the initial viewing. This analysis allowed us to assess whether changes in event structure explained memory above and beyond individual differences present at first viewing.

### Code and resource availability

The Python code used to reproduce all the results in this study is available at https://github.com/narjes-alzahli/EventTimescaleChanges. The results in MNI space can be accessed at https://neurovault.org/collections/21744/.

### Data availability

The analyses in this study used a publicly available dataset and stimuli, accessible at OpenNeuro: https://openneuro.org/datasets/ds001545/versions/1.1.1.

## Results

### Changes in Event Structure in The Brain with Repeated Movie Viewing

To identify changes in event structure in the brain, we examined TR-by-TR brain activity patterns during each of the six viewings of each movie clip (**Figs. 1-3**). To assess the strength of event structure at different timescales, we computed the similarity of spatial activity patterns within versus across HMM-defined event boundaries (Within vs. Between-Event Similarity) at slow and fast timescales (2 and 10 events) across the six viewings of each clip. We first conducted method validation, confirming that our approach of measuring Within vs. Between-Event Similarity provided reliable estimates of event structure across datasets/participants (see Methods/Within-vs. Between-Event Similarity and **Fig. S1**). Having shown that our measure is reliable, we next turned to examining our main question about *changes* in event structure. Change in event structure at a specific timescale (slow or fast) was defined as the difference between the average Within- vs. Between-Event Similarity values across viewings 2-6 and the Within- vs. Between-Event Similarity value during the first viewing.

Our analysis of the Intact clip revealed that the majority of brain regions (with reliable event structure for at least one timescale) showed no significant changes in event structure at either slow or fast timescales over movie repetitions (**Fig. 4**). Among regions that did show event structure changes across repeated viewings, the direction and magnitude of these shifts varied, such that we observed evidence for both finer event structure (weaker slow timescale structure and/or stronger fast timescale structure with repeated viewings) and coarser event structure (stronger slow timescale structure and/or weaker fast timescale structure with repeated viewings) across different brain regions. Regions showing finer event representations with repeated viewing included fusiform gyrus, which showed weaker event structure at a slow timescale over clip repetition (**Fig. 4, searchlight 1**), and temporoparietal junction, which showed stronger event structure at a fast timescale over clip repetition (**Fig. 4, searchlight 2**). Regions showing coarser event representations with repeated viewing included superior temporal sulcus, which, over clip repetition, showed stronger event structure at a slow timescale and weaker event structure at a fast timescale (**Fig. 4, searchlight 3**), and inferior frontal cortex, which, over clip repetition, showed stronger event structure at a slow timescale (**Fig. 4, searchlight 4**).

**Figure 4.**
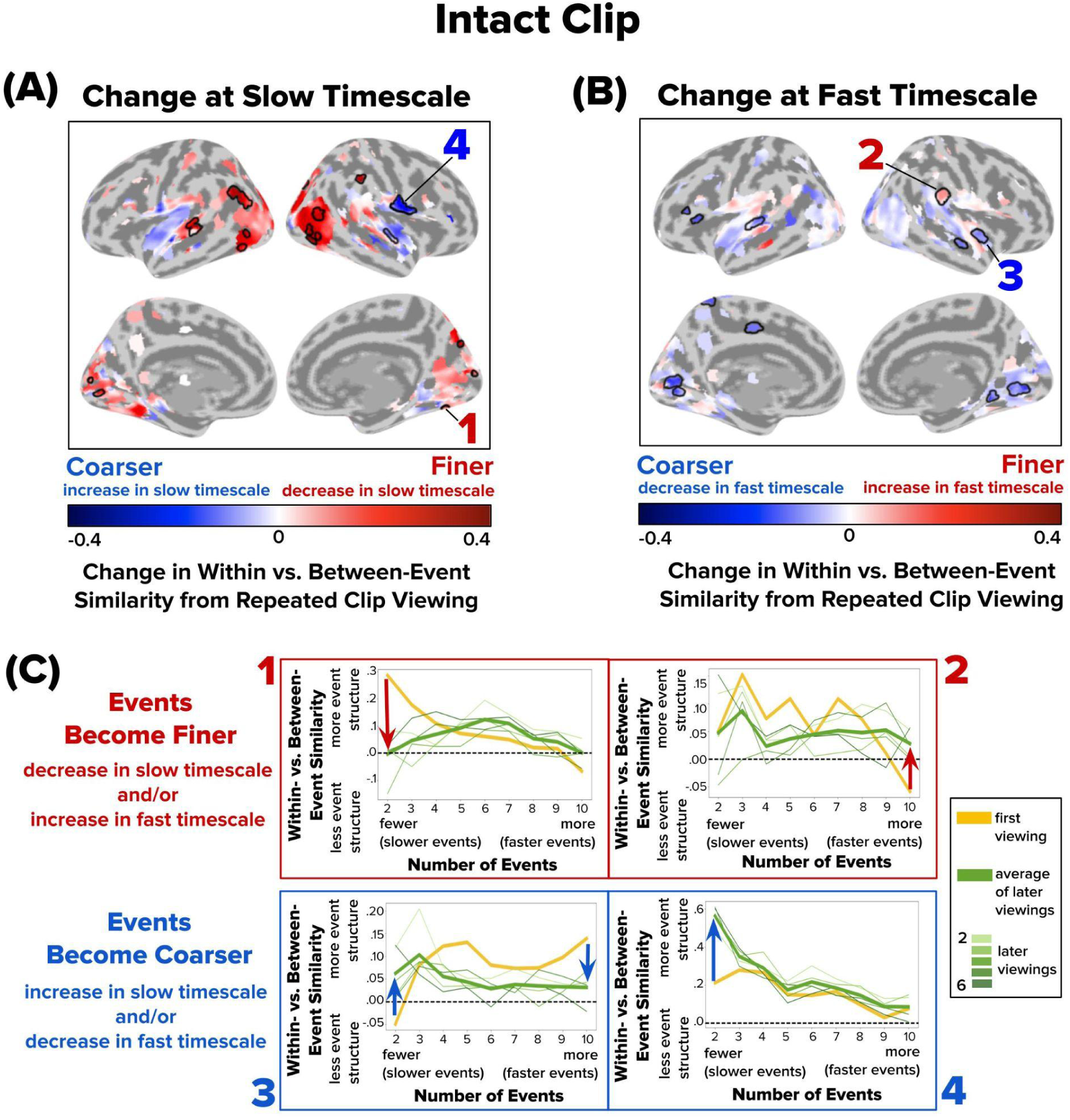
Changes in Event Structure at Slow and Fast Timescales with Repeated Viewing of the Intact Clip. Brain maps depict how repeated viewing of the Intact clip changes within- vs. between-event similarity at **(A)** a slow timescale (two events) and **(B)** a fast timescale (ten events). Values above zero indicate detectable event structure at that event count. The color gradient from blue to red indicates the direction of change from the initial viewing to subsequent viewings: blue represents a shift toward coarser/slower events, and red indicates a shift toward finer/faster events. Only searchlights with event structure for at least one timescale for each presentation are visualized. Significant searchlights are outlined with a black border; significance was determined using a permutation test with false discovery rate (FDR) correction (q < 0.05). Overall, events became finer in lateral visual cortex, and became coarser in ventral visual and lateral temporal cortex. **(C)** Four sample searchlights (1–4) were selected post hoc for illustration. Following the conventions in Figure 3, within-vs. between-event similarity at event counts 2-10 is shown for the first viewing (yellow line), subsequent viewings (thin green lines), and the average of subsequent viewings (thick green line). Red and blue arrows highlight the direction of change in within- vs. between-event similarity from the first viewing (thick yellow line) to the average of later viewings (thick green line).

We next repeated this analysis for the Scrambled-Fixed clip. Similarly to the Intact clip, the majority of brain regions with reliable event structure showed no significant changes in event structure across repetitions of the Scrambled-Fixed clip (**Fig. 5**). The few regions that showed significant shifts showed changes only at the slow timescale. Superior temporal gyrus displayed finer event structure with repeated viewing, by exhibiting weaker event structure at a slow timescale (**Fig. 5, searchlight 1**). Superior temporal sulcus displayed coarser event representations with repeated viewing by exhibiting stronger event structure at a slow timescale (**Fig. 5, searchlight 2**).

**Figure 5.**
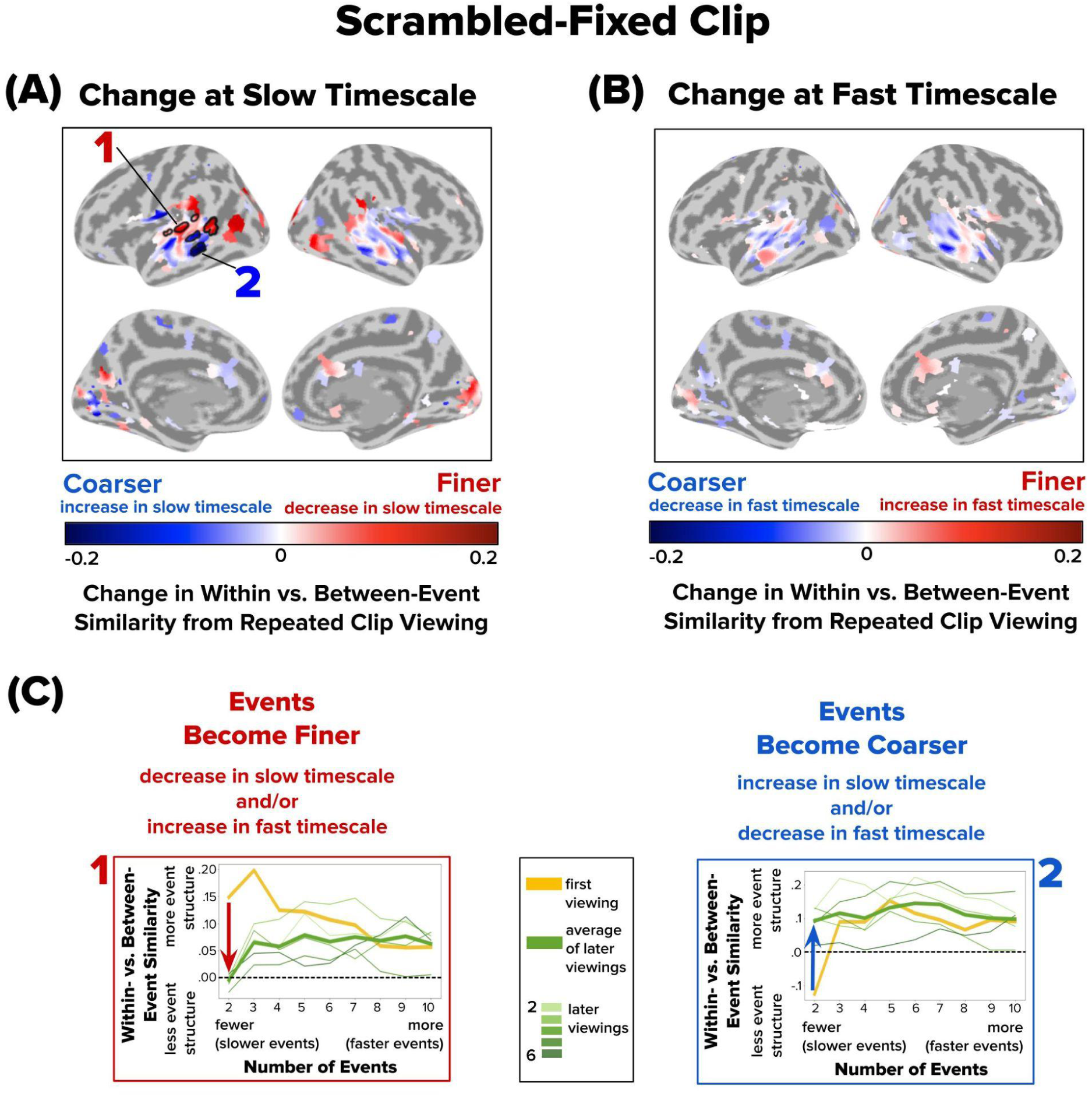
Changes in Event Structure at Slow and Fast Timescales with Repeated Viewing of the Scrambled-Fixed Clip. Following the conventions of Figure 4, brain maps depict how repeated viewing changes within- vs. between-event similarity at **(A)** a slow timescale (two events) and **(B)** a fast timescale (ten events). Significant searchlights are outlined with a black border (q < 0.05). Only subtle changes in timescales were observed for the Scrambled-Fixed condition, primarily in lateral temporal cortex. **(C)** Two sample searchlights (1–2) were selected post hoc for illustration. Red and blue arrows highlight the direction of change in within- vs. between-event similarity from the first viewing (thick yellow line) to the average of later viewings (thick green line).

Finally, we turned to the Scrambled-Random condition. As with previous clips, most brain regions with reliable event structure did not show significant changes in event structure across repetitions (**Fig. 6**). However, several regions exhibited significant shifts. Superior occipital cortex, cuneus, and collateral sulcus demonstrated finer event representations with repeated viewing, due to weaker slow-event structure in later viewings (**Fig. 6, searchlights 1 and 2**). Most regions, however, exhibited coarser event representations with repeated viewing, often due to stronger event structure at a slow timescale in later viewings. These regions included middle temporal gyrus and inferior frontal lobe, which exhibited both stronger slow-event structure and weaker fast-event structure in later viewings (**Fig. 6, searchlights 3 and 4**). Overall, for the Scrambled-Random clip, 39 out of 45 searchlights with significant timescale changes (87%) became coarser with repeated viewing, compared with 12/23 (52%) for the Intact clip and 1/3 (33%) for the Scrambled-Fixed clip.

**Figure 6.**
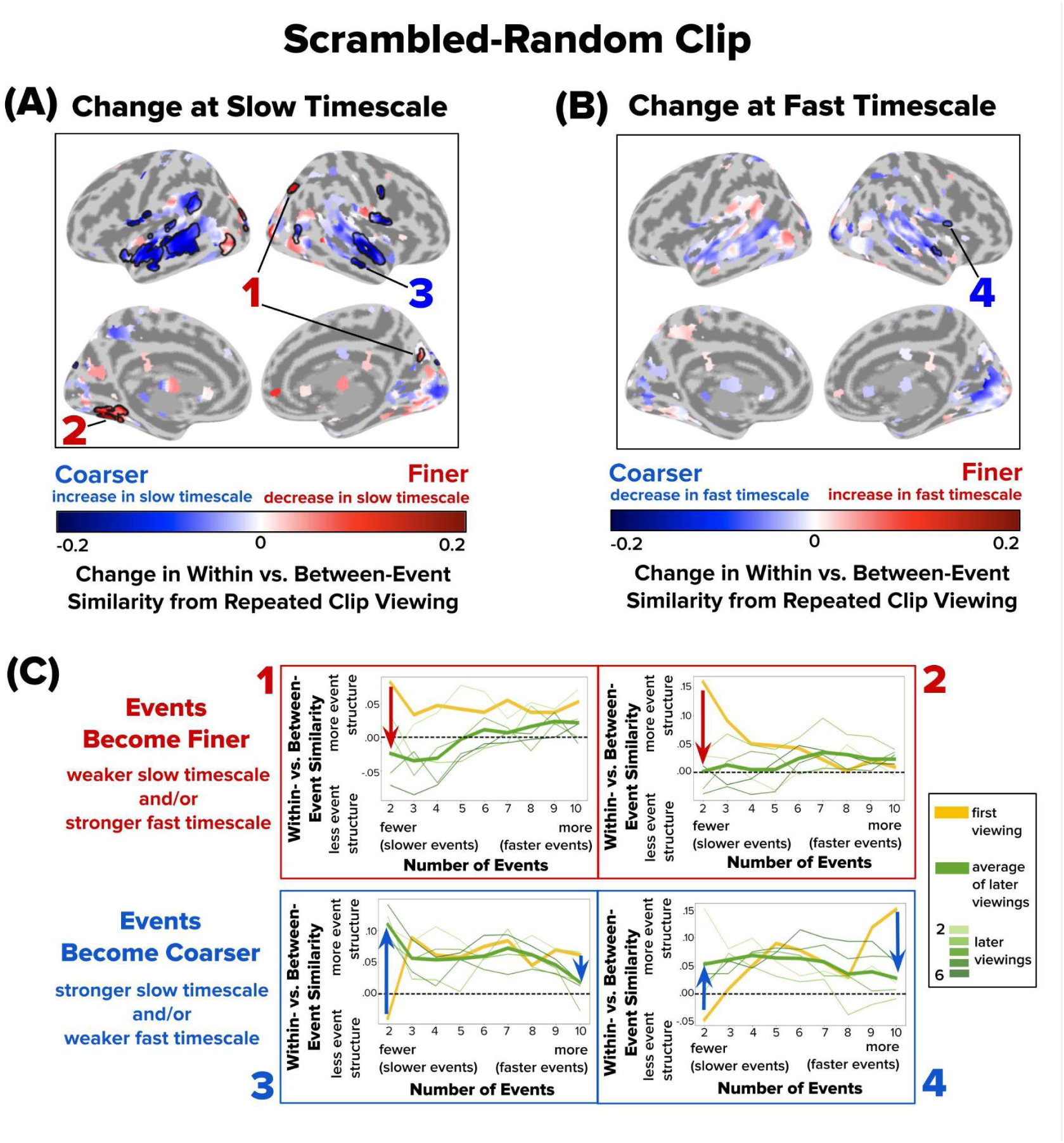
Changes in Event Structure at Slow and Fast Timescales with Repeated Viewing of the Scrambled-Random Clip. Following the conventions of Figures 4 and 5, brain maps depict how repeated viewing changes within- vs. between-event similarity at **(A)** a slow timescale (two events) and **(B)** a fast timescale (ten events). Significant searchlights are outlined with a black border (q < 0.05). Repeated viewing of the Scrambled-Random clip led to coarser event timescales throughout the lateral temporal cortex. **(C)** Four sample searchlights (1–4) were selected post hoc for illustration. Red and blue arrows highlight the direction of change in within- vs. between-event similarity from the first viewing (thick yellow line) to the average of later viewings (thick green line).

### Examining across-clip overlap in timescale changes

To identify brain regions that exhibited a consistent direction of change in event structure across different movie clips, we performed a conjunction analysis across the Intact, Scrambled-Fixed, and Scrambled-Random conditions (**Fig. 7**). This analysis does not assume shared event boundaries across clips; rather, it tests whether a region shows a consistent directional shift in slower or faster event structure with clip repetition, regardless of the specific sequence of events in any one clip. The analysis revealed changes in event structure that generalized across movie content. Some regions, such as lateral occipital cortex, showed a consistent weakening of event structure at the slow timescale in later viewings across all clips, reflecting finer event structure with repeated viewing (**Fig. 7, searchlights 1-2**). Other regions, across superior and middle temporal gyri, exhibited a consistent strengthening of event structure at the slow timescale in later viewings, reflecting coarser event structure over repetitions (**Fig. 7, searchlights 3-4**). These findings suggest that certain brain regions adapt the timescale of event segmentation in a systematic manner across varied stimulus contexts. Specifically, a consistent weakening of slow-timescale event structure in later viewings may allow a region to become more sensitive to fine-scale temporal details, whereas a consistent strengthening of slow-timescale structure in later viewings may support integration on longer temporal horizons. The consistency of these effects across move clips with different content may therefore index the general tendency of a brain region toward abstraction/integration vs fine-grained sensitivity. We return to this issue in the Discussion.

**Figure 7.**
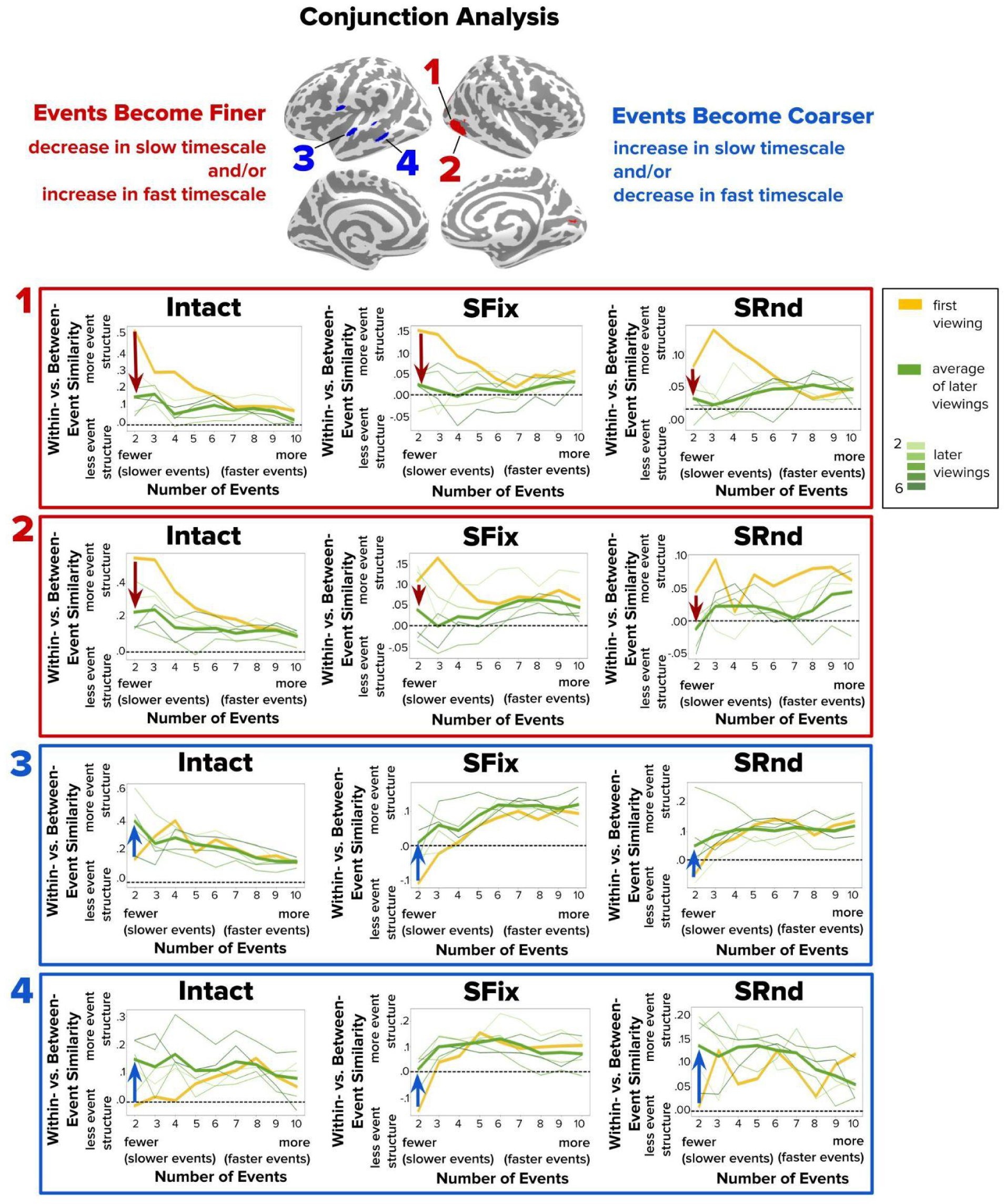
Conjunction Analysis Showing Consistent Changes in Event Structure Across All Clip Conditions (Intact, Scrambled-Fixed, Scrambled-Random). Brain regions from a conjunction analysis designed to identify areas showing significant changes in within- vs. between-event similarity across repeated viewings of all three clip conditions. Top two rows show sample searchlights for which events become finer with repetition across all clips, due to repetition-related weakening of event structure at the slow timescale of two events (downward red arrows). Bottom two rows show sample searchlights for which events become coarser with repetition across all clips, due to repetition-related strengthening of event structure at the slow timescale of two events (upward blue arrows). SFix = Scrambled-Fixed condition; SRnd = Scrambled-Random condition.

### Relating changes in event structure to event memory

We next aimed to determine the behavioral relevance of event structure changes in the brain by examining the correlation between these changes and subsequent memory. We focused on the six brain regions that showed a directionally consistent change in event structure across all three clips (as identified in the conjunction analysis) and tested if these changes predicted average recall across the clips in an individual differences analysis. Our initial analyses focused on only these regions to limit multiple comparisons. We report uncorrected p-values for this exploratory analysis but emphasize that future work with larger samples will be needed to confirm and generalize our findings.

We first examined overall memory performance by quantifying the number of details recalled for each clip (**Fig. 8**; see Materials and Methods/Scoring of event memory). As previously reported for this dataset in <u>Aly et al. (2018)</u>, memory for the Intact clip (M = 25.75, SD = 11.24) was better than memory for the Scrambled-Fixed clip (M = 21.45, SD = 12.62) (t_29_ = 2.17, p = .04, 95% CI: 0.24– 8.36) and the Scrambled-Random clip (M = 19.93, SD = 8.82) (t_29_ = 3.36, p = .002, 95% CI: 2.28–9.36). Recall scores for the Scrambled-Fixed and Scrambled-Random clips were not significantly different (t_29_ = 0.66, p = .51, 95% CI:−3.17–6.20).

**Figure 8.**
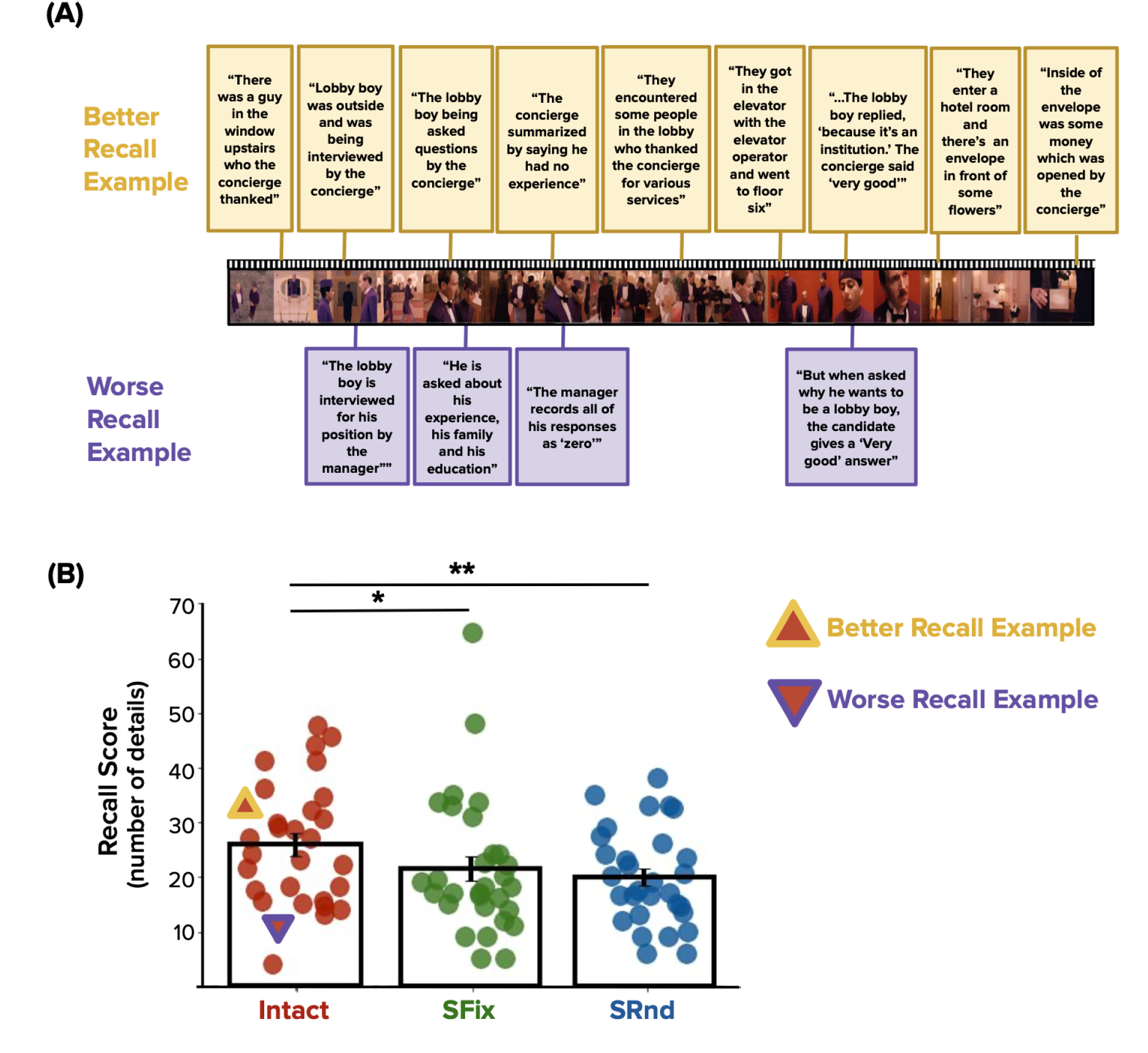
Memory Recall Examples. **(A)** Sample frames are shown from the Intact clip. Above and below these frames (connected by lines) are, respectively, examples of better and worse recall for the corresponding events. Responses from the worse recall example (bottom row, purple borders) are characterized by a relatively small number of vague, less detailed, or less accurate recollections of the events depicted in the clip. Responses from the better recall example (top row, yellow borders) are characterized by a relatively large number of detailed, accurate, and contextually rich recollections of the clip’s content. The worse recall example is shown in its entirety; only a subset of the better recall example is shown due to space constraints. **(B)** Recall scores for the Intact, Scrambled-Fixed (SFix), and Scrambled-Random (SRnd) conditions. Recall was superior for the Intact vs. Scrambled conditions. The better and worse recall examples for the Intact condition, shown in **(A)**, are highlighted in **(B)** with a yellow upward-facing triangle and purple downward-facing triangle, respectively. * p<0.05; ** p<0.01.

To relate changes in event structure to memory recall, we first estimated the change in Within- vs. Between-Event similarity values for each participant for a given searchlight at the timescale (slow or fast) that exhibited a significant group-level effect in that region. This was done for each clip separately and then averaged across clips to yield a single value for each participant. Finally, we obtained the correlation between this value and participants’ average recall across the three clips, separately for each searchlight.

Lateral occipital cortex (**Fig. 9A, left panel**, r=-0.324, p=0.0238, uncorrected) and middle temporal gyrus (**Fig. 9A, right panel**, r=-0.310, p=0.0341, uncorrected) showed correlations with recall performance. In both regions, participants with greater weakening of slow-timescale event structure had significantly better recall. Interestingly, this effect was observed despite contrasting group-level changes in these brain regions, with lateral occipital cortex exhibiting group-level weakening of slow timescale structure with clip repetition (**Fig. 7, searchlight 1**) and middle temporal gyrus exhibiting group-level strengthening of slow timescale structure with clip repetition (**Fig. 7, searchlight 4**).

**Figure 9.**
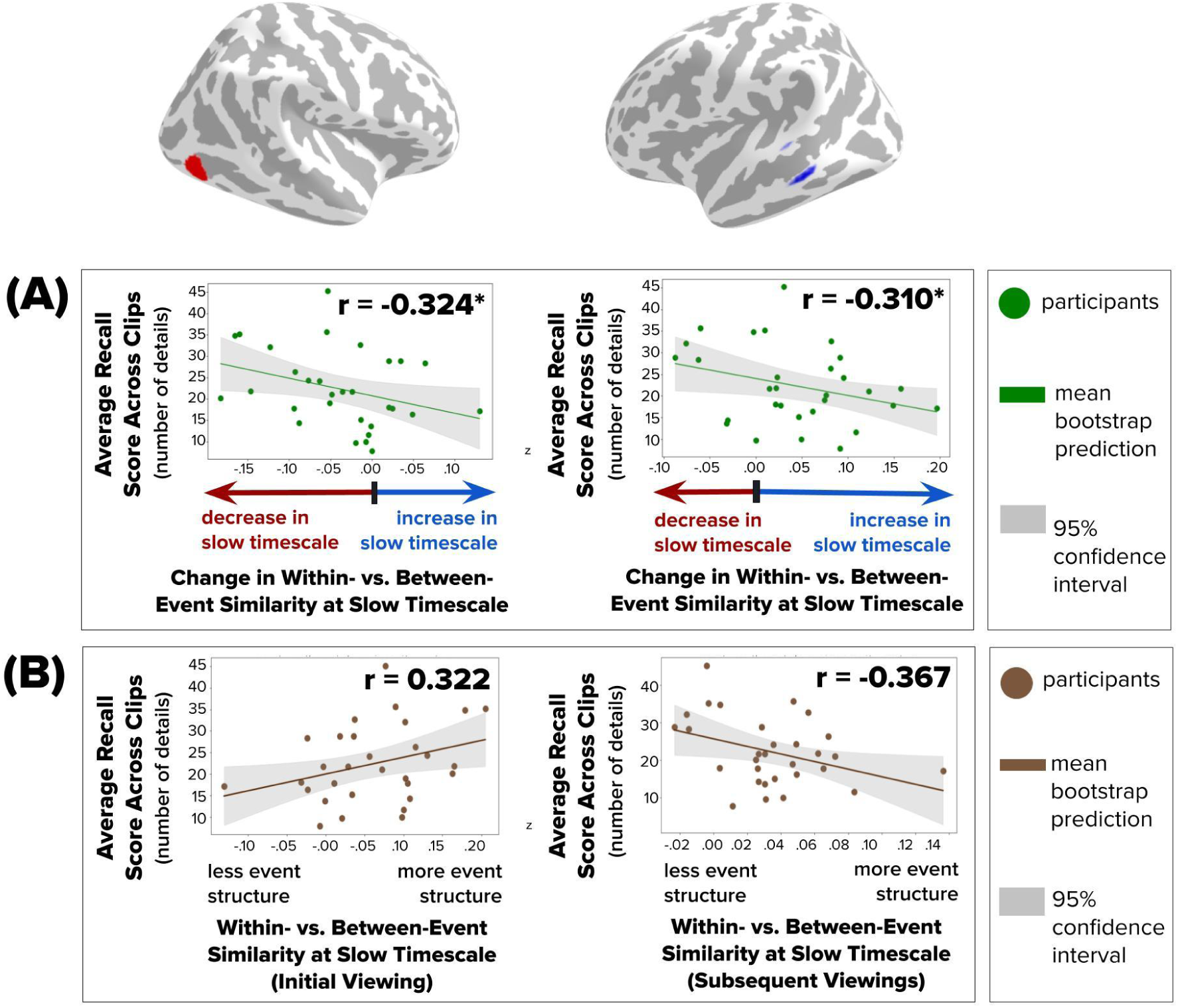
Across-Participant Correlation between Changes in Within- vs. Between-Event Similarity and Recall. For two brain regions identified via conjunction analysis across all three clip conditions (Intact, SFix, SRnd), each individual participant’s average recall score across all clips is plotted against their average change in event structure for 2 events across all clips. Left panels show a lateral occipital cortex region with group-level weakening of slow timescale structure with repetition of all three clips (red arrow at 2 events). Right panels show a middle temporal gyrus region with a group-level strengthening in slow timescale structure with repetition of all three clips (blue arrow at 2 events). For both regions, participants with greater weakening of slow-timescale structure had better recall (p < .05, uncorrected; exploratory).

To determine whether these correlations are driven by baseline differences between individuals at the first viewing or true change across repetitions, we conducted an analysis that predicted memory from the change in Within- vs. Between-Event Similarity with repeated viewing while controlling for baseline Within- vs. Between-Event Similarity from the initial viewing. Under this model, the effect in middle temporal gyrus remained significant (r=-0.347, p=0.0326, uncorrected), whereas the effect in lateral occipital cortex did not. To understand this pattern more directly, we then examined the pairwise relationships between memory and Within- vs. Between-Event Similarity from the initial viewing and, separately, the later viewings. This showed a clear difference: in lateral occipital cortex, slow-timescale event structure for the initial viewing positively predicted recall (**Fig. 9B, left panel**, r=0.322, p=0.0336, uncorrected), whereas in middle temporal gyrus, slow-timescale event structure for subsequent viewings negatively predicted recall (**Fig. 9B, right panel**, r=-0.367, p=0.0276, uncorrected). Thus, although both regions showed correlations with memory, the underlying sources of these relationships differ. These results suggest a potentially intriguing possibility that slow-timescale structure early in learning could be helpful for initial memory encoding, but may be a signature of poor detail memory if it is still present later in learning. An alternative possibility is that the different effects across regions arise because slow timescale structure best predicts memory whenever that structure is most prominent (initial viewings for LOC, subsequent viewings for MTG)—perhaps because of increased statistical sensitivity when the mean (and variance across participants) increases.. We will briefly consider these exploratory results further in the Discussion.

Finally, we asked whether there were additional brain–behavior relationships for individual clips. To do so, we ran another exploratory analysis that considered searchlights that showed either strengthening or weakening of event structure at either slow or fast timescales for any clip, and examined the correlation between those event structure changes and recall for that corresponding clip. After controlling for multiple comparisons across searchlights, no clip-specific correlations were significant.

## Discussion

### Summary of findings

We found both stability and flexibility in the brain’s event timescales. While many regions showed consistent event timescales across repeated viewings of a film clip, others exhibited changes. These included regions whose event representations became finer with repeated viewing, due to stronger event structure at a fast timescale in later viewings and/or weaker structure at a slow timescale in later viewings; and regions whose event representations became coarser, due to stronger event structure at a slow timescale in later viewings and/or weaker structure at a fast timescale in later viewings. The Intact clip—with preserved narrative coherence—elicited topographically organized timescale changes: in visual and auditory regions, such as lateral occipital cortex and middle temporal gyrus, repeated viewing led to finer event segmentation (weaker slow-timescale structure over clip repetition). In anterior regions involved in higher-order processing, such as inferior frontal cortex, event representations became coarser (stronger slow-timescale structure over clip repetition). This spatial organization of finer vs. coarser shifts was less apparent in the Scrambled-Fixed condition and largely absent in the Scrambled-Random condition, where coarsening was dominant. Thus, narrative coherence supports spatially organized timescale changes with increasing familiarity, whereas disorder in narrative structure prompts a consistent shift toward coarser event representations. Coarser representations for narratively incoherent events (the Scrambled conditions) may aid in integrating information over longer timescales—perhaps as the brain attempts to impose structure where little exists. Finally, the strength of coarse event structure was associated with recall in lateral occipital cortex and middle temporal gyrus. These results reveal that the brain’s event timescales can flexibly change with experience.

### Relation to prior work

Our findings build on evidence that the brain’s timescales are flexible and adjust to the stimulus statistics (<u>Lerner et al., 2014</u>; <u>Baumgarten et al., 2021</u>; <u>Çatal et al., 2024</u>). Dynamics in early auditory cortex and higher-order areas rescale in time to match compressed or dilated speech (<u>Lerner et al., 2014</u>). The timescale of neural responses lengthens and shortens with slowed and sped-up sensory information, respectively. Further, when predicting tone sequences played at half, normal, or double speed, the brain integrates a fixed amount of information—a constant number of tones—rather than a fixed time window (<u>Baumgarten et al., 2021</u>). Our observations align with such adaptive capacity, but extend it to repeated naturalistic viewing—rather than observing how timescales adapt to *external* sensory manipulations, we observe timescale changes due to increasing familiarity with a film clip, reflecting *internal* adaptation.

Our findings may be related to changes in stimulus predictability. According to a popular theory of event segmentation, event boundaries are triggered when incoming information deviates from predictions, forcing an update of the current “event model” (<u>Kurby & Zacks, 2009</u>). On initial viewing, many moments may be surprising, leading to frequent prediction errors and fine segmentation. With repetition, events become expected; a reduction in surprise could allow the brain to merge events into longer segments (<u>Hard et al., 2006</u>). Simultaneously, increasing knowledge of the plot might enable detection of subtle event transitions (e.g., minor details or foreshadowing). Under alternative theories of event segmentation, which argue for a primary role of contextual instability (rather than prediction error) in shaping event boundaries (<u>Güler et al.,</u> <u>2025</u>; <u>Shin & DuBrow, 2020</u>; <u>Shim et al., 2023</u>), this enhanced sensitivity to previously-ignored details could result in additional fine boundaries that were initially overlooked. Thus, experience can recalibrate the brain’s event structure at multiple scales.

The familiarity-dependent changes that we observed can also be understood within a hierarchical framework of event representation. Prior research suggests that event boundaries vary continuously in strength, such that strong boundaries mark major narrative transitions whereas weaker boundaries correspond to finer-grained sub-events (<u>Ben-Yakov & Henson, 2018</u>; <u>Lee & Chen, 2024</u>). Consistent with this, event boundaries may be organized in a partially nested hierarchy across the cortex, with finer states embedded within coarser states, rather than a single preferred segmentation (<u>Geerligs et al., 2022</u>). Under this view, multiple potential segmentations may coexist during perception, and which segmentation grain (fine or coarse) is more strongly expressed in a region may depend on factors such as predictability, attention, or prior knowledge. Our multiscale method makes this explicit by quantifying coarse and fine event structure simultaneously within a region, allowing us to observe how repeated experience alters which segmentation timescale dominates the neural signal. From this perspective, the fine- and coarse-timescale changes we observed do not necessarily imply that new events are created or old ones erased. Rather, familiarity may change the boundary “threshold” that determines which boundaries are more strongly expressed neurally: with repeated viewing, weak but pre-existing boundaries may become more salient (leading to stronger fine-timescale structure), or multiple fine transitions may be consolidated into a more stable macro-event (leading to stronger coarse-timescale structure). This interpretation aligns with work showing that sensitivity to boundary strength and the neural impact of boundaries can vary with learning, expectation, and memory demands (<u>Ben-Yakov & Henson, 2018</u>; <u>Lee & Chen, 2024</u>; <u>Lee et al., 2021</u>; <u>Barnett et al.,</u> <u>2024</u>; <u>De Soares et al., 2024</u>). Thus, our findings support a view in which hierarchical event representations are not rigidly dictated by fixed temporal receptive windows, but are dynamically reweighted with experience. Familiarity restructures the balance of nested timescales within a region, revealing that the hierarchy of event processing includes both stable architectural constraints and familiarity-driven changes.

This hierarchical framework also helps us interpret how temporal and narrative structure influence neural timescale changes across our stimulus conditions. For the Intact clip, storyline predictability and cohesion allowed the brain to fine-tune the narrative into meaningful sub-events with experience. In contrast, the Scrambled-Random condition, which lacked coherent structure, showed widespread coarsening in event segmentation with experience. This coarsening may be related to participants’ ability to gradually extract the narrative across repeated viewings, as suggested by their subsequent recall of the overall plot. However, due to the scrambled order of segments, shorter-term predictability was limited, and fine-tuning of event boundaries was less pronounced. The Scrambled-Fixed condition, with stable event order but an incohesive narrative, exhibited both coarsening and fine-tuning—suggestive of the brain balancing prediction of upcoming events with gradual extraction of narrative structure. However, relatively limited changes in event structure in this condition warrant caution in overinterpreting these results. Together, our results showcase the brain’s adaptive capacity to balance the extraction of global structure and encoding of detailed information. They underscore the dynamic nature of neural event segmentation and the importance of considering stimulus characteristics when interpreting neural timescale changes.

We speculate that changes in the brain’s event representations with experience may influence, or reflect, detection of larger-scale structure and/or meaningful sub-events. Yet, this flexibility poses a methodological challenge. Repetition of dynamic stimuli (e.g., <u>Huth et al., 2016</u>; <u>Golland et al., 2007</u>; <u>Hasson et al., 2009</u>) may qualitatively alter temporal event structure in neural responses. Thus, researchers should use caution when assuming that repeated presentations of dynamic stimuli leave the underlying neural event structure unchanged.

Our conjunction analysis aims to identify stimulus-agnostic, directionally consistent changes in event-structure strength at slow and fast timescales across distinct film clips. We interpret such cross-clip generalization as evidence that some regions adapt their integration windows via mechanisms not tied to any one clip’s boundary timings or content. Regions for which events consistently become coarser with repetition may have a tendency toward integration or abstraction, whereas regions for which events consistently become finer may exhibit increased sensitivity to detail as transitions grow familiar. Thus, the conjunction analysis identifies brain regions whose event structure adapts with experience in a context-general way—not because clips share boundaries or content, but because that region shows a general tendency toward abstraction vs. fine-tuning as events grow familiar . In contrast, regions whose timescale changes do not generalize across movie clips may be sensitive to properties that vary across the clips, such as their narrative coherence, predictability, or visual or semantic content. These context-sensitive regions may have tendencies toward abstraction or fine-tuning that are shaped by the amount of narrative cohesion (enabling abstraction) or temporal predictability (enabling fine-tuning) of a given clip.

Among regions showing cross-clip generalization, lateral occipital cortex and middle temporal gyrus were unique in also showing a relationship with recall performance. Prior work has shown that lateral occipital cortex supports object- and feature-based visual representations (<u>Grill-Spector & Kanwisher, 2001</u>). Because such representations capture rapidly changing visual details, familiarity may have enhanced sensitivity to these rapid visual transitions, sharpening fine-timescale segmentation in LOC over repeated viewings. Middle temporal gyrus, by contrast, has been linked to semantic and situational integration during language and narrative comprehension (<u>Binder et al., 2009</u>; <u>Leshinskaya & Thompson-Schill, 2020</u>). Building on this prior work, the strengthening of slow-timescale structure in this region may reflect increased abstraction of overarching narrative or situational context with increasing familiarity. The consistency of these effects across clips—despite their distinct narrative structures and temporal stability—suggests that lateral occipital cortex and middle temporal gyrus may contribute to segmentation at representational levels that go beyond specific storylines. Lateral occipital cortex may track low- to mid-level visual transitions that occur in all dynamic scenes, regardless of narrative context, whereas middle temporal gyrus may integrate conceptual or situational information over time in a way that abstracts beyond the surface structure of any one clip. The cross-clip consistency in timescale changes in these regions therefore likely reflects general properties of perceptual and semantic reorganization rather than sensitivity to particular boundary locations or plot coherence. Importantly, these patterns describe group-level tendencies in how regions adapt with experience, which may differ from how individual variation within those regions relates to behavior.

Considering individual differences, both lateral occipital cortex and middle temporal gyrus showed a relationship between timescale changes and recall performance, but our follow-up analyses indicate that these relationships arise from different underlying sources. In lateral occipital cortex (LOC), more slow-timescale structure during the initial viewing predicted more detailed recall. One possibility is that individuals who impose greater long-timescale structure on the incoming visual stream at first exposure encode more coherent perceptual representations, providing a strong, initial “scene scaffold” that later supports detailed recall (a scaffold that other downstream areas can potentially build on). The subsequent loss of slow-timescale structure in LOC, with clip repetition, does not appear to influence recall. . In middle temporal gyrus (MTG), even though the group average shows strengthening of slow event structure with clip repetition (as described above), individuals with less slow timescale structure during later viewings tend to recall more details. The relative lack of slow timescale structure may help preserve finer event boundaries and prevent collapsing information into gist, thereby preserving item- and moment-level details. As mentioned earlier, because these interpretations are post hoc, we emphasize that they should be viewed as tentative. Further, it is possible that the differential effects across LOC and MTG arise because each region shows a relationship with memory during the viewing in which slow timescale structure was most apparent in that region; this could, in turn, reflect greater statistical sensitivity to individual differences when the mean (and variance) in slow timescale event structure are higher. Future work is needed to arrive at a more definitive understanding of why these regions relate to memory, and whether effects may vary for memory assessed via recall of details vs. gist.

### Limitations and alternative explanations

We interpret our results as qualitative changes in the structure of neural event representations with experience, but might they be due to other factors? One concern is regression to the mean: regions with initially strong event structure at a particular timescale might appear to shift toward intermediate values on later viewings. However, some regions with initially strong structure at the slow timescale exhibited an additional increase in this structure with repeated viewing, inconsistent with regression to the mean (e.g., searchlight 4 in **Fig. 4**). Further, many regions showed stable temporal structure with repeated viewing, which would be unexpected if our results were a statistical artifact. Finally, we only considered regions with statistically reliable event structure for each clip viewing, reducing the risk of observing changes due purely to noise.

Another concern that viewers might become bored during later clip presentations, and increasing disengagement may drive changes in neural event structure. We think this is unlikely to explain our results. First, disengagement should lead to weakening of both fine and coarse event structure with repeated viewing: idiosyncratic mind wandering or distraction should produce widespread loss of consistency in event representations across people, but our analyses require similar event representations across individuals. Further, we observed *selective* changes in event structure with movie repetition: changes were not widespread across all timescales, they were not widespread across the brain, they varied by stimulus type, and the direction of changes reversed across brain regions and across fast and slow timescales within a region.

A potential limitation also arises from our memory measure. We used the number of recalled details as an index of memory, which could align with finer event segmentation (parsing events more finely might produce more event fragments to describe). Though widely used in narrative recall research (<u>Martinez, 2024</u>; <u>Chen et al., 2017</u>; <u>Silva et al., 2019</u>), future studies should include complementary metrics (e.g., recall coherence; recognition memory) to test whether the relationship between neural event timescales and memory differs for gist memory vs. detail memory.

Finally, although we repeated identical stimuli multiple times, this rarely occurs in natural settings—primarily when we rewatch movies or listen to the same song again and again. Even when stimuli are identical across repetitions, contextual factors (goals, affect, setting) can modulate how event timescales adapt. We view our design as a tractable proxy for examining the effects of recurring structure on event representations in the brain. Future work will be important for determining the generalizability of our findings to event structure like that found in daily routines (e.g., commute routes, morning rituals), and stimuli that repeat in variable contexts.

### Conclusion and future directions

Brain regions’ event representations are flexible and can be reshaped by experience. Regions dynamically adjust how they segment continuous information, either compressing or expanding event durations as events become familiar. Neural timescales are therefore not static properties of each region—they can adapt to the observer’s knowledge.

Future research can leverage our new method, which quantifies event structure at multiple timescales simultaneously within a brain region, rather than assigning it a single preferred timescale. Tracking how both slow and fast event structures evolve *within a brain region* with repeated experience allows researchers to characterize flexible, multi-level segmentation dynamics across the brain. Additionally, future studies can explore how changes in event timescales relate to the perception of elapsed time (<u>Matthews, 2011</u>; <u>Eaglement, 2010; Rose</u> <u>& Summers, 1995</u>; <u>Sherman & Yousif, 2025</u>)—for example, whether strengthening of slow-timescale structure with repeated viewing in higher-order regions is associated with the perception of time slowing with repetition. Finally, do timescale changes persist, or does the brain revert to its initial segmentation patterns after a delay? Does persistence of timescale changes predict durable memory, whereas reversion to initial segmentation patterns predict forgetting? Addressing these questions will elucidate how the brain flexibly constructs and updates event representations, and how this flexibility supports learning, re-interpreting, and remembering real-world experiences.

Together, our findings lay important groundwork for studying the brain’s representation of events, revealing that even the temporal “building blocks” of experience, events themselves, can reorganize with experience. This opens the door to understanding how the brain’s dynamic balancing of stability and flexibility supports our ability to understand and remember complex narratives.

## Acknowledgements

This work was funded by a National Institutes of Health Research Project Grant (R01EY034436 to M.A. and C.B). We thank the Alyssano Group for valuable feedback on this project.

## Acknowledgements

This work was funded by a National Institutes of Health Research Project Grant (R01EY034436-01) to M.A. and C.B. We would like to thank the Alyssano Group for helpful advice on this project.

## Author Contributions

**Narjes Al-Zahli**: Conceptualization, Data Curation, Formal analysis, Methodology, Project Administration, Software, Validation, Visualization, Writing—original draft, Writing—review and editing; **Mariam Aly**: Conceptualization, Methodology, Funding Acquisition, Resources, Project Administration, Supervision, Writing—original draft, Writing—review and editing; **Chris Baldassano:** Conceptualization, Methodology, Funding Acquisition, Resources, Project Administration, Supervision, Writing—original draft, Writing—review and editing.

## Supplementary Figures

**Figure S1.**
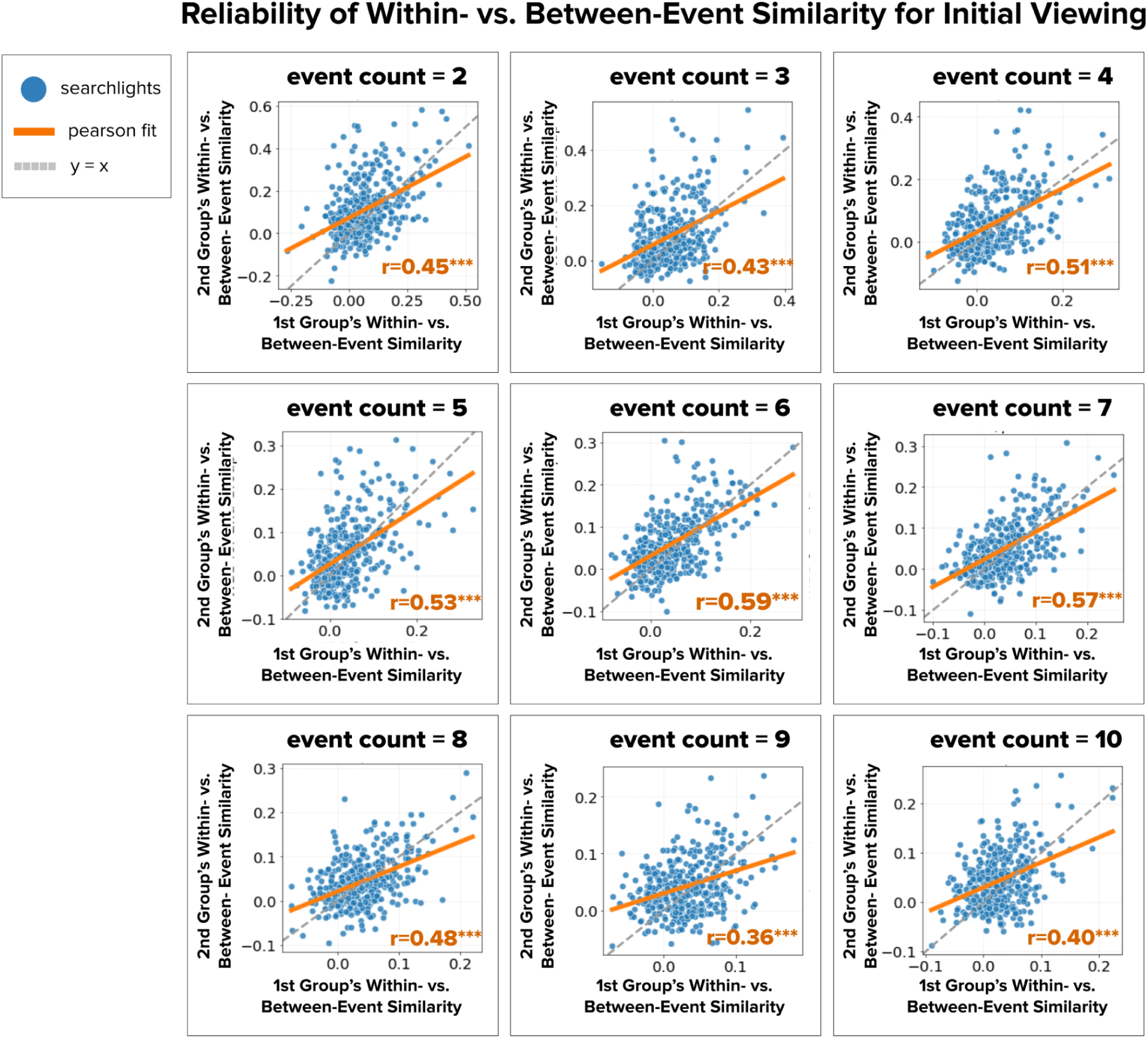
Validation of Within- vs. Between-Event Similarity (Intact Clip). To assess the reliability of the Within- vs. Between-Event Similarity measure during participants’ initial viewing of the movie clip, we examined how correlated this measure was across searchlights for two independent participant samples. Participants were first randomly divided into two independent groups of 15 participants each. Then, within each group separately, participants were randomly divided into training and testing subsets. We identified HMM event boundaries for each searchlight in the training subset and then used those boundaries to compute Within- vs. Between-Event Similarity in the testing subset. Training and testing subsets were then swapped, and results were averaged across five different participant splits within each independent group. Finally, we examined the correlation in Within- vs. Between-Event Similarity values across the two independent groups. Reliability for Within- vs. Between-Event Similarity values was consistently high across all event counts, demonstrating that this measure is relatively stable across subsets of participants.

**Figure S2.**
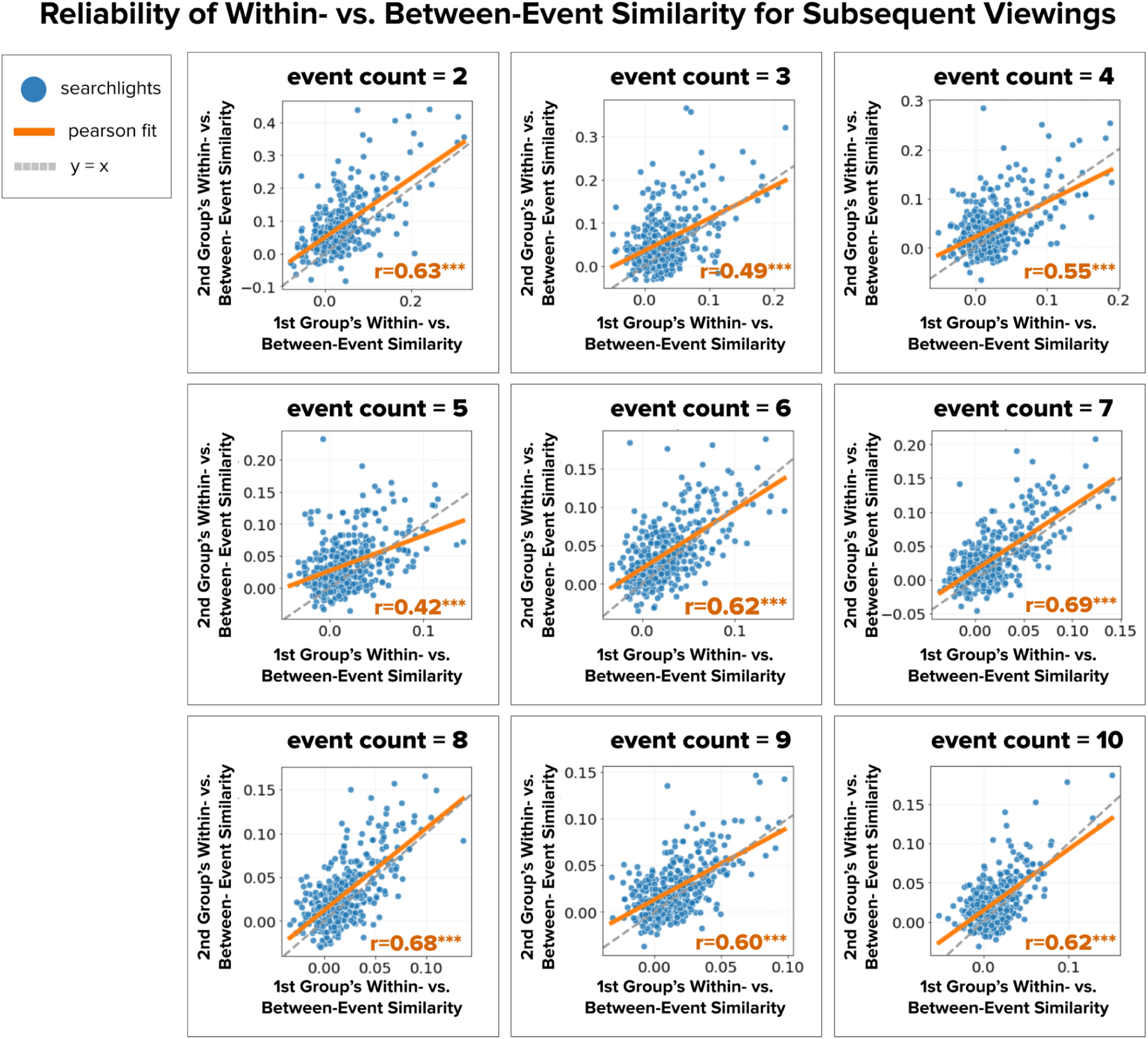
Validation for Subsequent Viewings (Intact Clip). The same split-half reliability procedure was repeated for participants’ subsequent viewings of the film clip. Results again showed strong reliability across all event counts, indicating that the Within- vs. Between-Event Similarity measure remains robust across repeated viewings.

**Figure S3.**
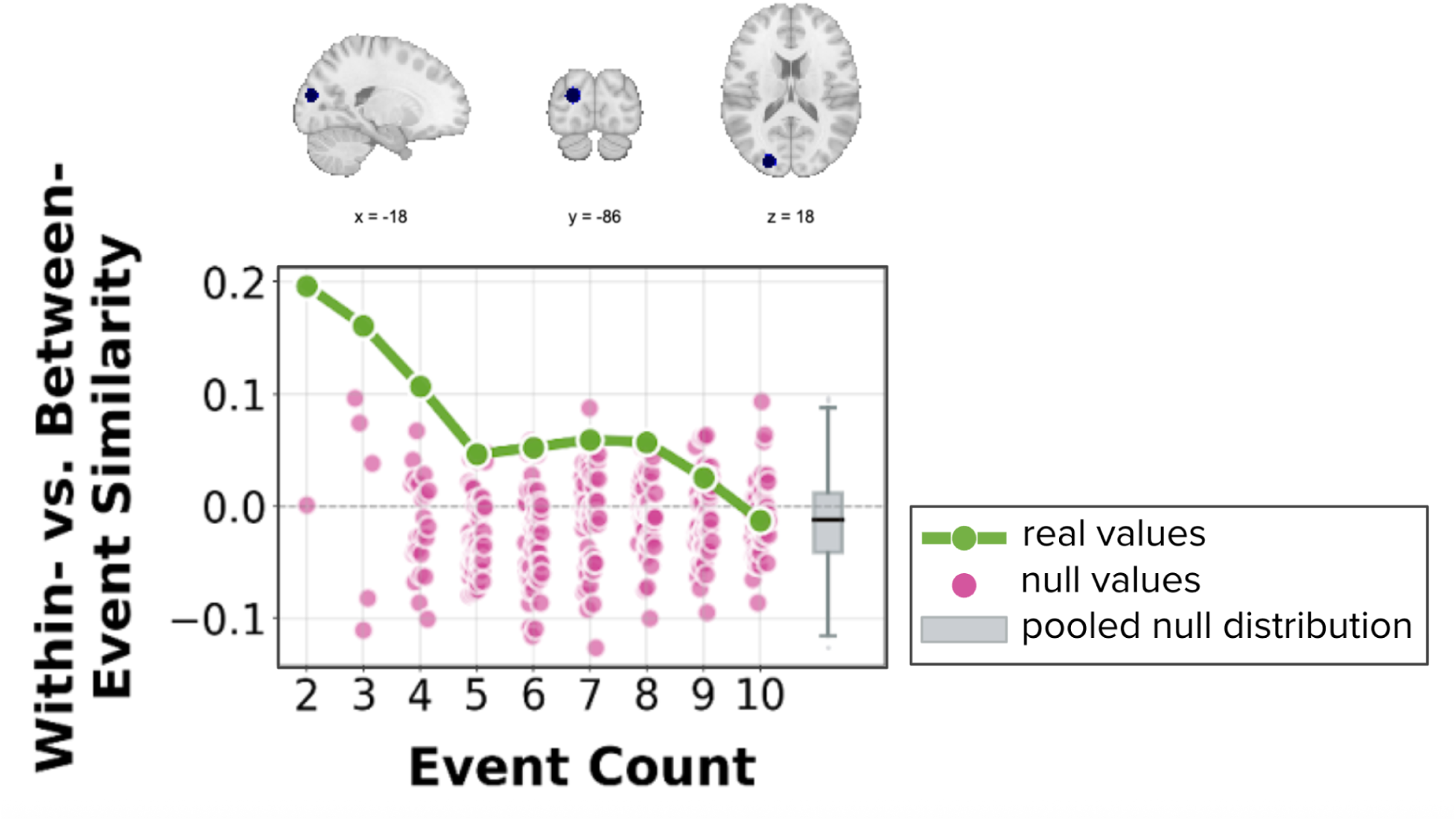
Within- vs. Between-Event Similarity Values for Observed Data and Null Distributions. Shown is a sample searchlight in left occipital cortex (data from the fourth viewing of the Intact clip). The figure plots Within- vs. Between-Event Similarity as a function of the number of events for the observed data (green line). Pink dots show Within- vs. Between-Event Similarity when event segment lengths were randomly shuffled for each event count, producing per-event-count null distributions.The gray boxplot shows the pooled null distribution across event counts. Null distributions show no systematic shifts across event counts, indicating that pooling across event counts does not introduce bias. This searchlight was included in the main timescale analysis and was selected post hoc for illustrative purposes.

**Figure S4.**
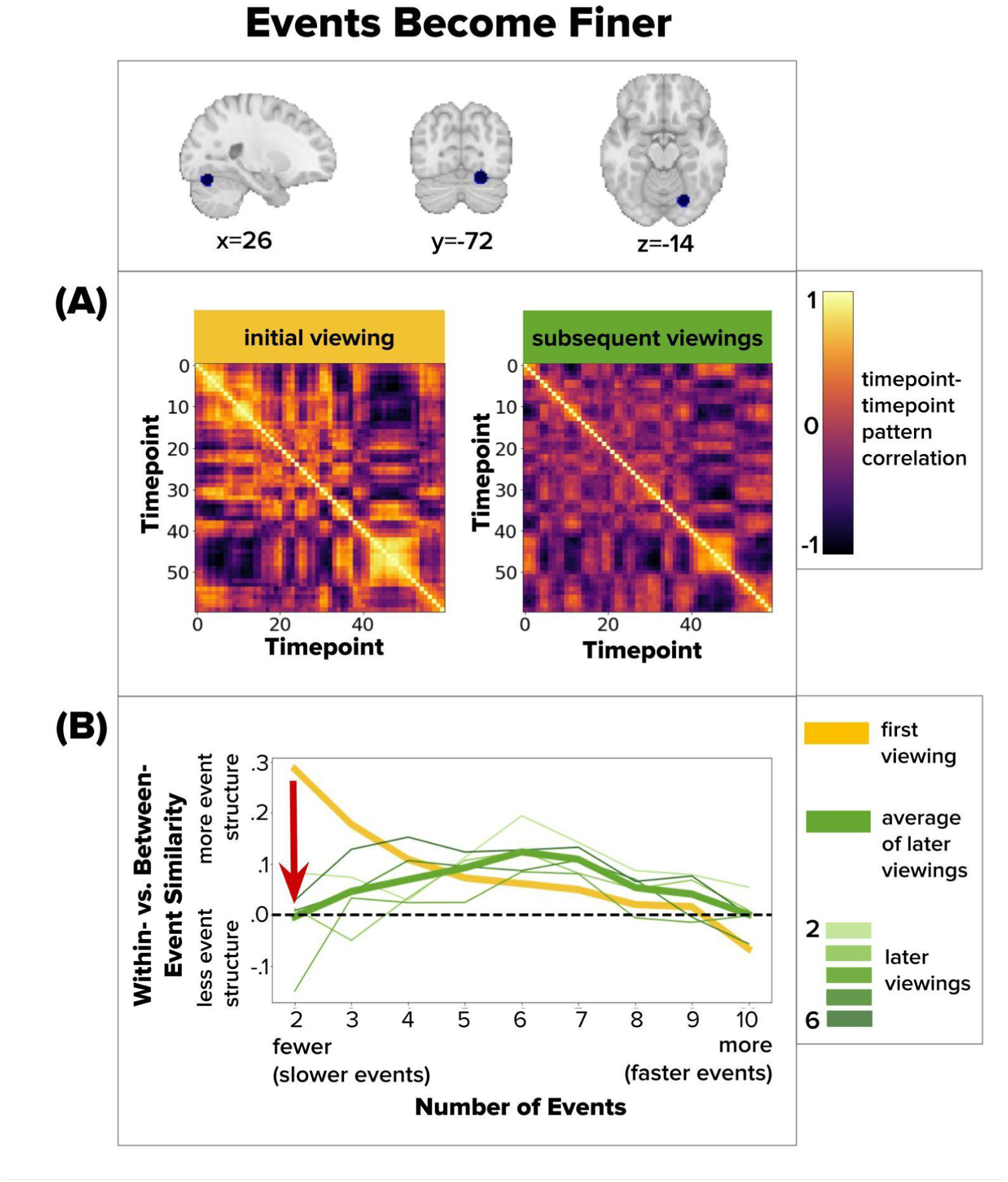
Timepoint-by-timepoint similarity matrices illustrate a significant decrease in slow-timescale event structure with repeated viewing of the Intact clip. **(A)** Timepoint-by-timepoint pattern-similarity matrices from initial viewing (left) and the average of subsequent viewings (right). Color intensity reflects the similarity between fMRI spatial activity patterns at each pair of timepoints. Timepoints that are highly similar to one another tend to be grouped into the same neural event. During the first viewing, the matrix shows stronger slow-timescale structure, visible as fewer, larger high-similarity blocks along the diagonal. During subsequent viewings, this slow-timescale structure diminishes, replaced by more, smaller blocks, indicating a shift toward finer-grained event segmentation. **(B)** Following the conventions of Figures 4–6, the red arrow indicates the direction of change in Within- vs. Between-Event Similarity from the first viewing (thick yellow line) to the average of later viewings (thick green line), reflecting a decrease in slow-timescale event structure over repetitions. The figure shows a searchlight in fusiform gyrus that showed a significant timescale change for the Intact clip. It was selected post hoc for illustration.

**Figure S5.**
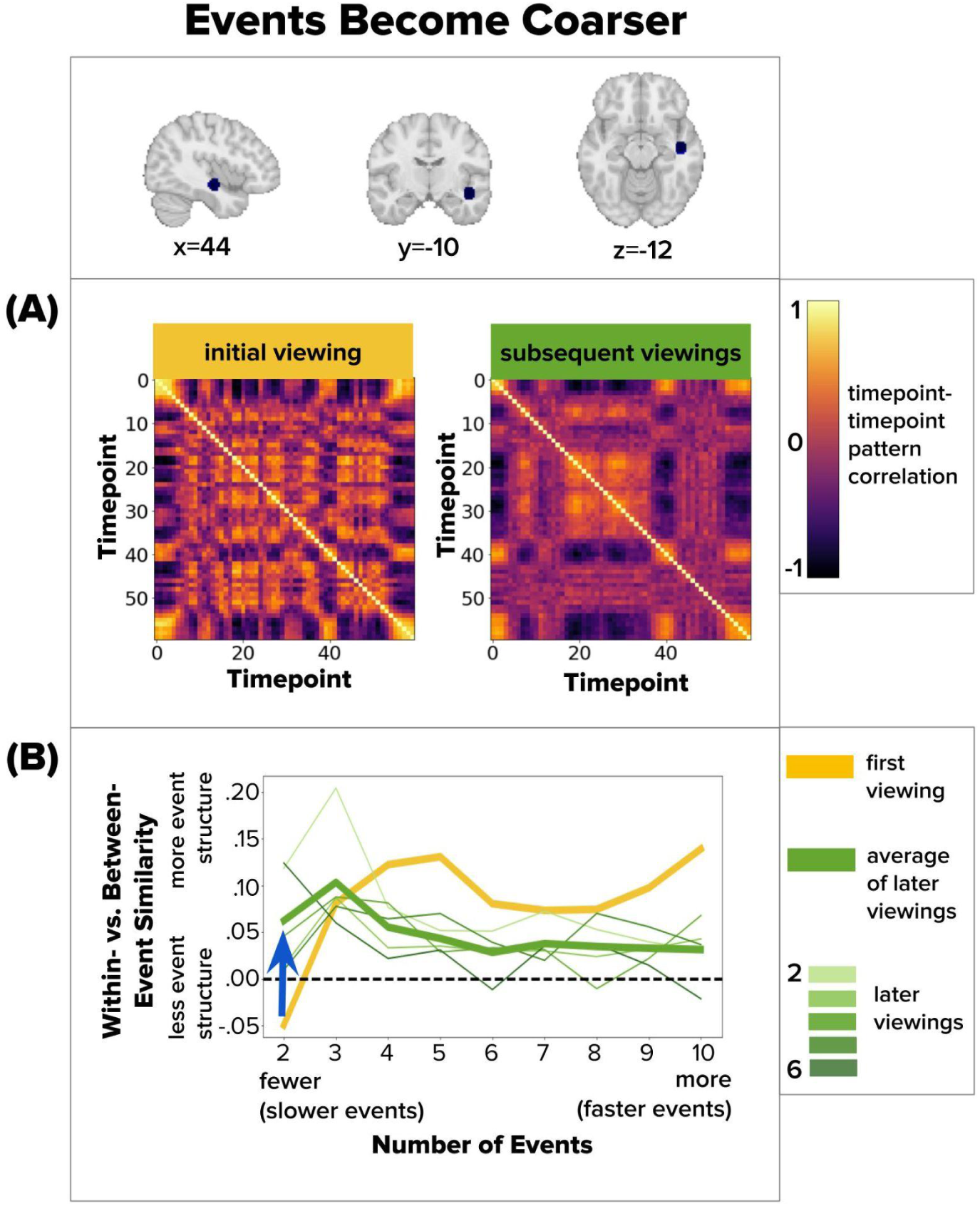
Timepoint-by-timepoint similarity matrices illustrate a significant increase in slow-timescale event structure with repeated viewing of the Intact clip. **(A)** Following the conventions of Figure S4, slow-timescale event structure appears stronger with repeated viewing: larger, more coherent blocks of high within-event similarity are evident. **(B)** Following the conventions of Figure 4-6, the blue arrow indicates the increase in within- vs. between-event similarity from the first to later viewings at the slow timescale. The figure shows a searchlight in superior temporal gyrus that showed a significant timescale change for the Intact clip. It was selected post hoc for illustration.

